# Module-Selection Balance in the Evolution of Modular Organisms

**DOI:** 10.64898/2026.04.01.715873

**Authors:** Mark Kim, Sarah M. Ardell, Sergey Kryazhimskiy

## Abstract

Fitness of an organism depends on different combinations of its traits in different environments. Therefore, how a population traverses the trait space during an adaptive walk in the current environment determines its fitness in future conditions. These trait dynamics depend on two fundamental features of the genotype-phenotype-fitness map (GPFM), its degree of pleiotropy and the entropy of the trait space (i.e., how densely different regions of the trait space are populated by genotypes), but these dependencies are not well understood. To address this gap, here we consider four simple two-trait GPFM models that differ in pleiotropy and trait-space entropy. We find that on a GPFM with universal pleiotropy and a high-entropy fitness optimum, populations evolve along the fitness gradient, which implies that the trait under stronger selection improves exponentially faster than the trait under weaker selection. At the opposite end of the GPFM continuum, on a variationally modular GPFM with a low-entropy fitness optimum, populations approach a quasi-steady state that we term a “module-selection balance” where both traits improve at proportional rates and their ratio remains constant. We find that convergence to a module-selection balance is robust with respect to several other GPFM properties, is enhanced by clonal interference and weakened by sex. Extrapolating from our simple models, we hypothesize that variationally modular organisms exhibit bi-phasic dynamics of genome evolution, especially in the clonal interference regime, and we find support for this hypothesis in the metagenomic data from Lenski’s long-term evolution experiment in the bacterium *Escherichia coli*. Our work outlines how the structure of pleiotropy and trait-space entropy constrain trait evolution. In particular, it suggests that the effective trait-space dimensionality of adapting variationally modular organisms may be much smaller than the total number of traits relevant for fitness.

## 1 Introduction

Heritable variation arises and is transmitted through generations at the genetic level, but natural selection acts on it only if this variation has a phenotypic manifestation. This duality implies that the efficiency of natural selection and the resulting dynamics of adaptation depend on the architecture of the genotype-phenotype-fitness map (GPFM). A salient feature of GPFMs of biological organisms is that they are modular, with several different types of modularity having been distinguished (Wagner et al., 2007). Most obviously, organisms are functionally modular, meaning that different physiological functions are performed by distinct units, such as protein complexes, metabolic pathways or organs (Hartwell et al., 1999; Ravasz et al., 2002; Bhattacharyya et al., 2006; Kreimer et al., 2008). Many studies have also found that a typical mutation affects only a small subset of key traits or functions (Cheverud et al., 1997; Dudley et al., 2005; Wagner et al., 2008; Wang et al., 2010; Wagner and Zhang, 2011; Miller et al., 2014; Collet et al., 2018; Kinsler et al., 2020, 2024; Donyavi et al., 2025; Ghosh et al., 2026). We refer to this property as “variational modularity” (Wagner et al., 2007; Wagner and Zhang, 2011) but note that this term is used somewhat differently in quantitative genetics (see Section 1 in the Supplementary Information for an extended discussion). Although many functional and variational modules coincide (e.g., each protein is encoded by a gene), this is not the case in general (e.g., different organs are not encoded by different parts of the genome). The ubiquity of modular GPFM architectures in biological systems raises two related but distinct questions (Wagner et al., 2007; Paaby and Rockman, 2013; Melo et al., 2016; Houle and Rossoni, 2022). Why and how do such architectures evolve? And how does variational modularity constrain the system’s evolutionary dynamics at the trait level?

Much of the previous literature addressed the former question, with a particular focus on understanding the conditions when various types of modularity evolve by natural selection. The emerging consensus is that modularity can increase the system’s evolvability and thereby be favored by “second-order” natural selection (reviewed by Wagner et al., 2007; Melo et al., 2016; Houle and Rossoni, 2022). In particular, certain kinds of modularity can evolve if the organism’s fitness depends on multiple functions that share some sub-functions (Lipson et al., 2002; Kashtan and Alon, 2005; Kashtan et al., 2007; Sun and Deem, 2007; Parter et al., 2008; Kashtan et al., 2009a,b; He et al., 2009; Espinosa-Soto and Wagner, 2010; Tikhonov et al., 2020) or when modularity enables the organism to preferentially generate variation along phenotypic directions that are most favored by selection (Wagner, 1996; Welch and Waxman, 2003; Pavlicev et al., 2010; Melo and Marroig, 2015; Chebib and Guillaume, 2022; Tikhonov et al., 2020; do O and Whitlock, 2023).

The question of how variational modularity constrains trait evolution has so far received less attention (but see relevant discussions by Arnold et al., 2001; Welch and Waxman, 2003; Schmiegelt and Krug, 2014; Park et al., 2015; Melo et al., 2016). Much of our current understanding of trait evolution is based on Fisher’s geometric model (FGM, reviewed by Tenaillon, 2014) and the related Simpson-Lande “adaptive landscape” model (reviewed by Arnold et al., 2001). Crucially, these models typically assume that mutations move the organism in random directions in the trait space, such that almost every mutation affects all traits, an assumption that is referred to as “universal pleiotropy” (Wagner and Zhang, 2011). However, as mentioned above, data suggest that organisms are variationally modular, in the sense that mutations tend to move the organism along certain preferred directions in the trait space whereas other directions may be inaccessible by any single mutation.

In general, biases in the supply of mutations can affect evolutionary dynamics (Yampolsky and Stoltzfus, 2001; Stoltzfus and McCandlish, 2017; Storz et al., 2019; Gomez et al., 2020; Cano and Payne, 2020; Cano et al., 2022; Sane et al., 2023; Tuffaha et al., 2023; Gitschlag et al., 2023; Parveen et al., 2025; Fuqua and Vakirlis, 2026; Barber and Couce, 2026). Moreover, a recent experimental study by Venkataram et al. (2020) suggests that variational modularity in particular can impose important constraints on these dynamics in asexual organisms. In this study, the authors evolved strains of *Escherichia coli* with defects in the translation machinery and found that adaptive mutations in this module were often outcompeted by mutations in other modules, which stalled improvements in translation. The authors termed this effect “evolutionary stalling”. Subsequent theoretical work by Gomez et al. (2020) showed that evolutionary stalling can occur in the so-called clonal interference regime (Schiffels et al., 2011; Good et al., 2012; Lang et al., 2013) whenever adaptive mutations that improve the stalled module are much less common and/or provide much smaller fitness benefits than mutations in other modules. These studies therefore suggest that models with universal pleiotropy may not adequately describe the dynamics of trait evolution of modular organisms. Thus, we set out to better understand these dynamics, with a particular focus on how typical adaptive trajectories in the trait space differ between GPFMs with variationally modular versus universally pleiotropic architectures.

To address this question, we consider a set of simple models that share the same phenotype-fitness map but differ in the genetic architecture of the underlying traits. Specifically, we consider fitness to be a concave unimodal function of two underlying functional traits, with a single combination of trait values maximizing fitness, as in the classic Fisher’s geometric model (Fisher, 1930; Tenaillon, 2014) and related models (Wagner, 1989; Arnold et al., 2001; Jones et al., 2003; Melo and Marroig, 2015; Chebib and Guillaume, 2022). We compare and contrast the evolutionary dynamics that are induced in this trait space by two distinct genotype-phenotype map architectures, one with universal pleiotropy and one with variational modules (Figure 1). In both architectures, we consider a genome with two pieces of DNA, which we refer to as “chromosomes”. In the GPFM with universal pleiotropy, each locus on each chromosome contributes to both functional traits, whereas on the modular GPFM, loci on chromosome 1 contribute only to trait 1 and loci on chromosome 2 contribute only to trait 2 (Welch and Waxman, 2003; Gutenkunst and Sethna, 2007).

**Figure 1.**
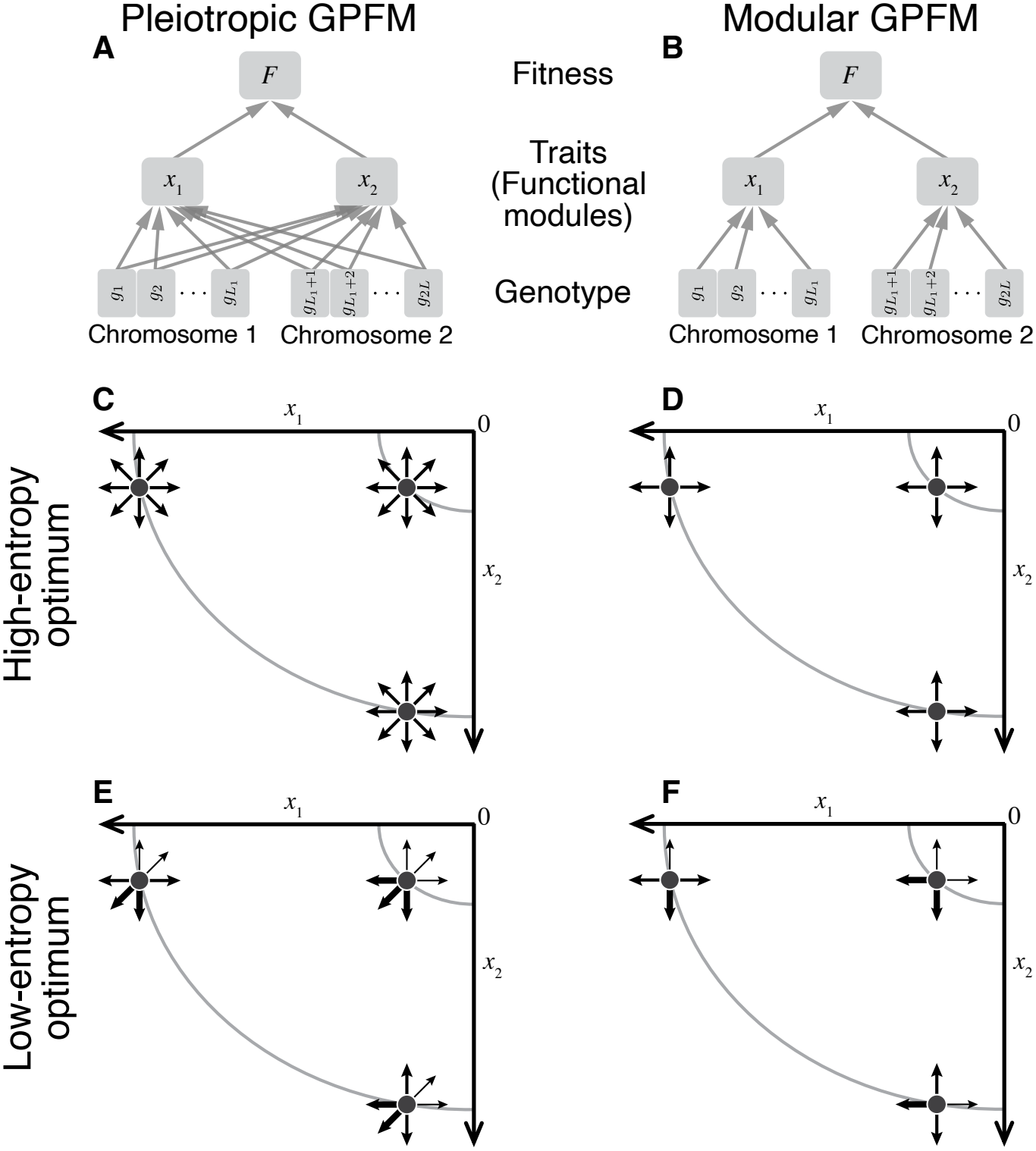
Schematic of the simple GPFMs models that differ by variational modularity and the entropy of the optimum. **A**. Pleiotropic GPFM. *F* is fitness; *x*_*i*_ is the value of quantitative trait *i* (which we think of as the performance of functional module *i*); *g*_*l*_ is the allele (0 or 1) at site _*l*_. Arrows indicate dependencies (see Section 2.1 for details). **B**. Variationally modular GPFM. Notations are the same as in A. **C**. Phenotypic effects of mutations in the pleiotropic GPFM with a high-entropy optimum. Each black circle represent a genotype and arrows represent mutations. Gray arcs show fitness isoclines. On this GPFM, the distribution of phenotypic effects of mutations changes little across genotypes located in the genotype-dense vicinity of the fitness optimum (Section 2.1.2). **D**. Phenotypic effects of mutations in the modular GPFM with a high-entropy optimum. Notations are as in panel C. On this GPFM, mutations on chromosome *i* affect only trait *i*, and the distribution of phenotypic effects of mutations changes little across genotypes located in the genotype-dense vicinity of the fitness optimum (Section 2.1.4). **E**. Phenotypic effects of mutations in the pleiotropic GPFM with a low-entropy optimum. Thickness of arrows indicates the supply of mutations. Other notations are the same as in panel C. On this GPFM, trait-improving mutations become increasingly rare in genotypes that are closer to the optima (Section 2.1.3). **F**. Phenotypic effects of mutations in the modular GPFM with a low-entropy optimum. Notations are the same as in panels C,E. On this GPFM, trait-improving mutations become increasingly rare in genotypes that are closer to the optima (Section 2.1.5).

Previous studies have found that the entropy of the trait space—that is, how densely this space is populated by genotypes—can also affect evolutionary dynamics (Iwasa, 1988; Sella and Hirsh, 2005; Schaper and Louis, 2014; Manrubia et al., 2021). In general, since some trait values are realized by many more genotypes than others (Schaper and Louis, 2014; Kinney, 2026), random mutations are biased toward producing these more common, high-entropy trait values. To probe the effects of entropy on the dynamics of trait evolution within the context of both pleiotropic and modular GPFMs, we consider two extreme cases with respect to the distribution of genotypes in the trait space, (i) when the vicinity of the optimal phenotype is densely populated by genotypes (“high-entropy optimum”) and (ii) when the vicinity of the optimal phenotype is genotype-sparse (“low-entropy optimum”).

In addition to the intrinsic properties of the GPFM, evolutionary dynamics depend on the mode of reproduction (sexual versus asexual) and on the supply of adaptive mutations (i.e., whether the population evolves in the successive versus concurrent mutations regime (Desai and Fisher, 2007)). In particular, when reproduction is asexual, both chromosomes are always inherited together (i.e., they are in complete genetic linkage with each other), whereas when reproduction is obligately sexual, the two sets of parental chromosomes are reassorted at every reproduction event. To understand how populations traverse the trait space during adaptation on different GPFMs in different evolutionary regimes, we use simulations and supplement them with analytical approximations whenever possible. Note that we model reassortment between chromosomes, but ignore recombination within chromosomes. We find that trait evolutionary trajectories change qualitatively depending on the architecture of the underlying GPFM and the entropy of the optimum, and the evolutionary regime is important as well. Most interestingly, we find that populations evolving on modular GPFMs converge to a quasi-stable state which we refer to as the “module-selection” balance, with this convergence being most pronounced when the optimum is low-entropy or when populations are asexual and evolve in the concurrent mutations regime. Finally, using metagenomic data from Lenski’s long-term evolution experiment, we show that *E. coli* in this experiment display signs of convergence to a module-selection balance.

Our approach is a blend of approaches practiced within population and quantitative genetics. Most previous studies of the FGM have been carried out in the framework of population genetics, focusing primarily on the fitness effects of mutations. In contrast, we focus on traits, which is the traditional purview of quantitative genetics. This focus is important because how fitness gains in the current environment manifest at the trait level can have major implications for the organism’s survival in future environments. For example, if a microbe adapts to the current nutrient-rich environment by losing its ability to sense certain nutrients (as is often observed in evolution experiments (Quan et al., 2012; Kvitek and Sherlock, 2013; Venkataram et al., 2016; Wu et al., 2020)), it may not be able to survive the next period of famine (Kvitek and Sherlock, 2013; Li et al., 2018), whereas equivalent fitness gains achieved by, say, improvements in translation are likely to be advantageous across many environments. In other words, different trait combinations that provide the same fitness and are therefore indistinguishable to selection in the current condition may be critically different in future conditions. While our approach is similar to quantitative genetics in focus, the mathematical apparatus that we use is that of population genetics. This is because standard quantitative genetics models abstract away the genetic architecture of traits, but modeling this architecture explicitly is necessary to understand how the supply of adaptive genetic variation changes across the trait space.

## 2 Methods

### 2.1 Genotype-phenotype-fitness map models

We consider a phenotype-fitness map that is based on Fisher’s geometric model (FGM) with *M* = 2 traits that contribute to fitness (Fisher, 1930; Tenaillon, 2014) (see Section 2.1.1 below). We construct the full genotype-phenotype-fitness map by coupling this phenotype-fitness model with 4+1 different genotype-phenotype map (GPM) models (see Sections 2.1.2–2.1.6). In all of our models, a genome with 2*L* bi-allelic loci, with alleles 0 and 1, is either implied or modeled explicitly, as specified below. In models where the genotypes are modeled explicitly, we consider two chromosomes, with each chromosome *i* containing *L*_*i*_ loci, such that *L*_1_ + *L*_2_ = 2*L*, and we consider only mutations that flip the bit at a single site. See Table 1 for the list of key parameters. Thus, GPFMs have the same two-chromosome organization but differ in how mutations affect the two traits. The four “simple” GPFMs, which are at the center of our investigation (Sections 2.1.2–2.1.5), all have additive genotype-phenotype maps, but also consider one GPFM to probe the consequences of deviations from additivity (see Section 2.1.6 below).

**Table 1.**
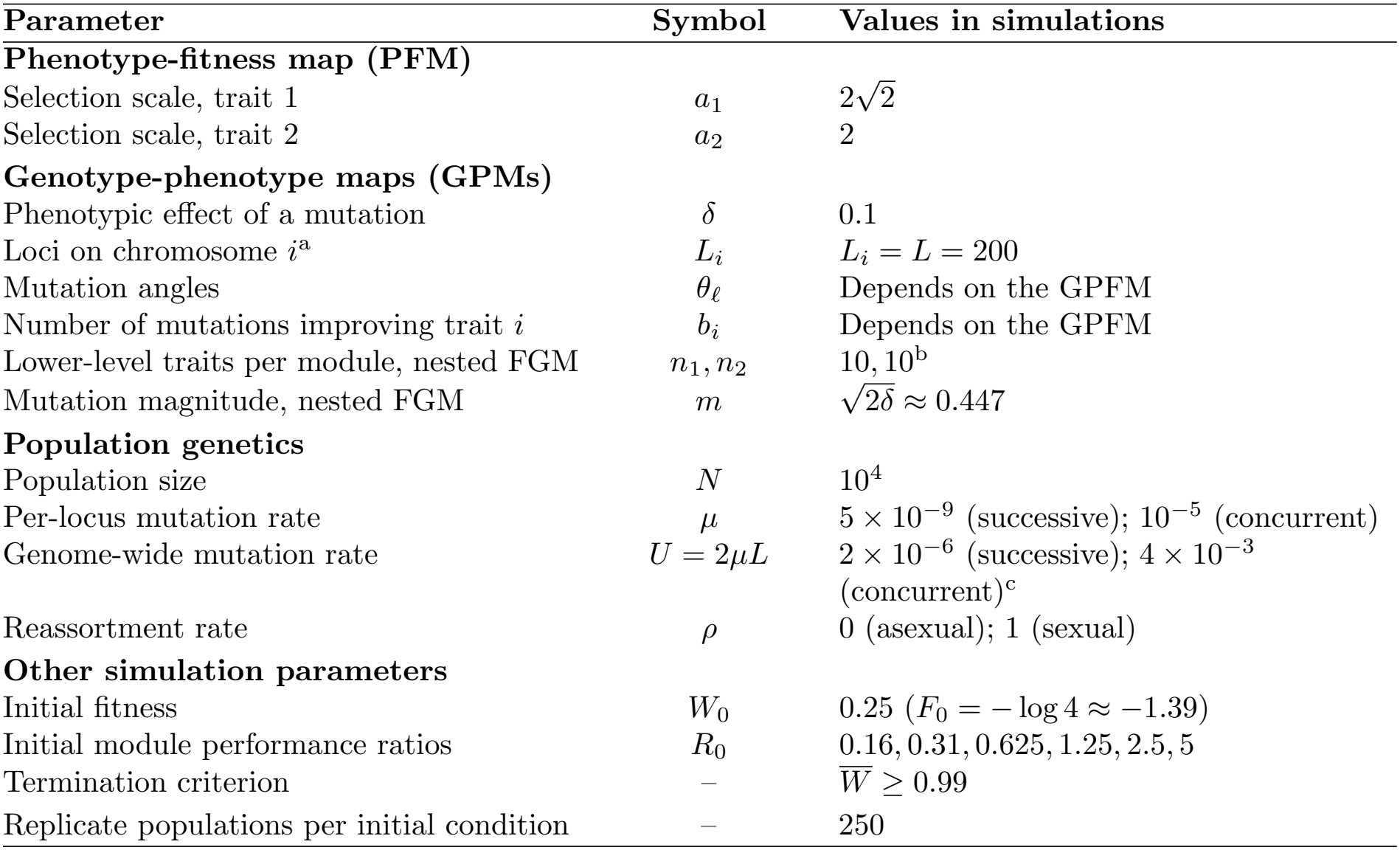
Key parameters and values used in the two-module simulations. Values apply to all models/simulations unless stated otherwise. ^a^The nested FGM is simulated without explicit loci, so *L* does not enter there, but both modules are assumed to be encoded by the same number of loci. ^b^Figure S5 shows results for *n*_1_ = 10, *n*_2_ = 20. ^c^In nFGM, *U* = 10^*−*7^ (successive) and *U* = 2 × 10^*−*4^ (concurrent), split equally between modules.

| **Parameter** | **Symbol** | **Values in simulations** |
|---|---|---|
| **Phenotype-fitness map (PFM)** |  |  |
| Selection scale, trait 1 | *a*<sub>1</sub> | 2√2 |
| Selection scale, trait 2 | *a*<sub>2</sub> | 2 |
| **Genotype-phenotype maps (GPMs)** |  |  |
| Phenotypic effect of a mutation | *δ* | 0.1 |
| Loci on chromosome *i*<sup>a</sup> | *L*<sub>*i*</sub> | *L*<sub>*i*</sub> = *L* = 200 |
| Mutation angles | *θ*<sub>*ℓ*</sub> | Depends on the GPFM |
| Number of mutations improving trait *i* | *b*<sub>*i*</sub> | Depends on the GPFM |
| Lower-level traits per module, nested FGM | *n*<sub>1</sub>, *n*<sub>2</sub> | 10, 10<sup>b</sup> |
| Mutation magnitude, nested FGM | *m* | $\sqrt{2\delta} \approx 0.447$ |
| **Population genetics** |  |  |
| Population size | *N* | 10<sup>4</sup> |
| Per-locus mutation rate | *μ* | 5 × 10<sup>−9</sup> (successive); 10<sup>−5</sup> (concurrent) |
| Genome-wide mutation rate | *U* = 2*μL* | 2 × 10<sup>−6</sup> (successive); 4 × 10<sup>−3</sup> (concurrent)<sup>c</sup> |
| Reassortment rate | *ρ* | 0 (asexual); 1 (sexual) |
| **Other simulation parameters** |  |  |
| Initial fitness | *W*<sub>0</sub> | 0.25 (*F*<sub>0</sub> = −log 4 ≈ −1.39) |
| Initial module performance ratios | *R*<sub>0</sub> | 0.16, 0.31, 0.625, 1.25, 2.5, 5 |
| Termination criterion | – | $\overline{W} \geq 0.99$ |
| Replicate populations per initial condition | – | 250 |

#### 2.1.1 Phenotype-fitness map

Our model of the phenotype-fitness map with two traits *x*_*i*_, *i* = 1, 2 is illustrated in Figure 1A,B. We can think of each trait *x*_*i*_ as representing the performance of a functional module *i* = 1, 2 relative to its optimum, so that *x*_*i*_ = 0 corresponds to peak performance in the current environment. Thus, we refer to *x*_*i*_ interchangeably as either *trait* or *module performance*. We denote the position of an individual (or the position of the population mean) in the two-dimensional trait space by the trait vector ***x*** = (*x*_1_, *x*_2_) (Figure 1C–F). If *W* is the Wrightian fitness, then we define the Malthusian fitness *F* = log *W* as a quadratic function of the trait vector ***x***,

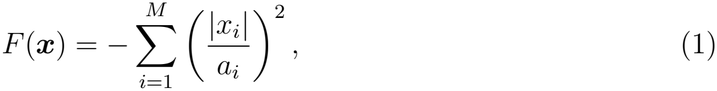

where the constants *a*_*i*_ determine the contribution of trait *i* to organismal fitness and hence the strength of selection on the corresponding module, with smaller *a*_*i*_ indicating stronger selection. Without loss of generality, we assume that *a*_2_ *< a*_1_, i.e., trait 2 is under stronger selection than trait 1. We initialize both traits with negative values (*x*_*i*_ *<* 0), so that adaptation corresponds to increases in *x*_*i*_ toward zero.

#### 2.1.2 Pleiotropic GPM with a high-entropy optimum

Our baseline genotype-phenotype map is a GPM with universal pleiotropy, where each locus contributes to both traits, and the high-entropy optimum, as shown in Figure 1A,C (see also the work by Blanquart et al. (2014); Hwang et al. (2017); Park et al. (2020) for related models that extend the canonical FGM to include an explicit genotype-phenotype map). Specifically, we define the traits of genotype *g* = *g*_1_*g*_2_ … *g*_2*L*_, with *g*_*l*_ ∈ {0, 1}, as

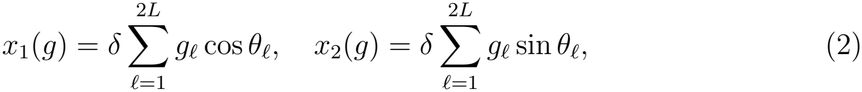

where *δ >* 0 is the magnitude of mutational effects and the angles *θ*_*l*_ are drawn independently from a uniform distribution on [0, 2*π*) (Figure 1C). The strictly optimal phenotype is encoded by a single genotype *g*_0_ = 00 … 0. However, by the central limit theorem, when *L* is large, the majority of genotypes are located within a circle of radius 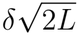 of the optimum.

Mutations that flip a bit at a single site _*l*_ displace the trait vector by vector ±***δ***_*l*_ = ±(*δ* cos *θ*_*l*_, *δ* sin *θ*_*l*_). In contrast to the canonical FGM, in our model, the number of mutational directions available to any genotype is finite. Therefore, the number of available beneficial mutations and their trait effects vary across the trait space not only because of the geometry of the phenotype-fitness map but also because of how the genotypes are distributed across the trait space, which we refer to as trait-space entropy (Iwasa, 1988; Sella and Hirsh, 2005; Schaper and Louis, 2014; Kinney, 2026). However, in large genomes (*L* ≫ 1) and with uniformly distributed *θ*_*l*_, mutations are approximately isotropic at sufficiently populated trait values—in this case, in the 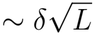 vicinity of the optimum—and our pleiotropic GPFM becomes asymptotically equivalent to the canonical FGM. Our analytical results are derived in this regime. Specifically, the fitness effect of a mutation with effect ***δ*** = (*δ* cos *θ, δ* sin *θ*) that occurs in a genetic background with trait vector ***x*** is

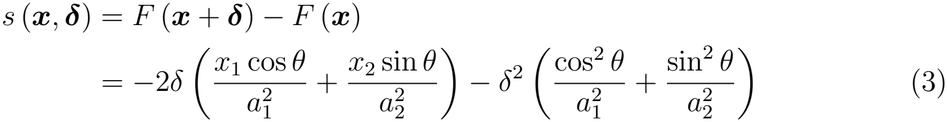

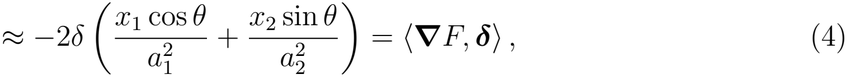

where 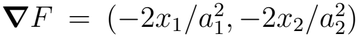 is the fitness gradient and ⟨***a, b***⟩ denotes the inner product of vectors ***a*** and ***b***.

#### 2.1.3 Pleiotropic GPM with a low-entropy optimum

We also consider a highly pleiotropic GPM with a low-entropy optimum. We construct such GPM by using the same expressions (2) for the traits of genotype *g* = *g*_1_*g*_2_ … *g*_2*L*_ (*g*_*l*_ ∈ {0, 1} ) but with the angles *θ*_*l*_ being drawn independently from a uniform distribution on [*π*, 3*π/*2] (Figure 1E). The strictly optimal phenotype is again encoded by a single genotype *g*_0_ = 00 … 0, but in this case, when *L* ≫ 1, the vast majority of genotypes are located within 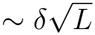 of the mean trait vector −2*δL/π***1**, where **1** = (1, 1), i.e., far from the optimum.

As in the high-entropy case, mutations that flip a bit at a single site _*l*_ displace the traits by vector ±***δ***_*l*_ = ± (*δ* cos *θ*_*l*_, *δ* sin *θ*_*l*_), and their selection coefficients are given by expressions (3), (4). In this model, all mutations that flip 1 → 0 are beneficial and all mutations that flip 0 → 1 are deleterious. Importantly, since the phenotype space is sparsely populated by genotypes near the phenotypic optima, beneficial mutations cannot be assumed to have angles that are uniformly distributed on [*π*, 3*π/*2]. For example, any genotype with coordinates with |*x*_2_| ≲ *δ* must have 1s only at sites with *θ*_*l*_ ≈ *π*. Because of this non-uniformity, the standard FGM approximation is not applicable and more advanced approaches must be used to obtain analytical results in this model, which we do not pursue here.

#### 2.1.4 Modular GPM with a high-entropy optimum

We model a GPM with variational modularity by assuming that loci on chromosome 1 contribute only to trait 1 and loci on chromosome 2 contribute only to trait 2 (Figure 1B). Specifically, the trait values of genotype *g* = *g*_1_*g*_2_ … *g*_2*L*_ are given by expressions (2) but with *θ*_*l*_ taking values 0 or *π* with equal probabilities for 1 ≤ _*l*_ ≤ *L*_1_ and *θ*_*l*_ taking values *π/*2 or 3*π/*2 with equal probabilities for *L*_1_ *<* _*l*_ ≤ 2*L*. Genotype 00 … 0 encodes the phenotypic optimum, but so do many other genotypes.

All mutations have the phenotypic effects of magnitude *δ*. By the central limit theorem, the number of loci carrying allele 1 among genotypes near the optimum is approximately normal with mean *L/*2 and variance *L/*4. Genotypes with trait values 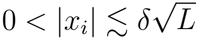 have access to approximately *b*_*i*_ = *L*_*i*_*/*2 beneficial mutations, i.e., approximately half of mutations are beneficial and half are deleterious. A beneficial mutation affecting module *i* has the selection coefficient

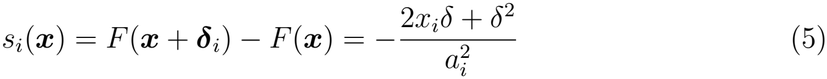

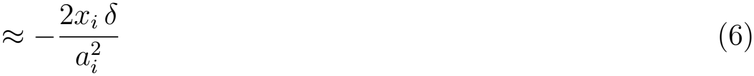

where the last approximate equality holds as long as *δ* ≪ |*x*_*i*_|.

#### 2.1.5 Modular GPM with a low-entropy optimum

The modular GPM with a low-entropy optimum is modeled analogously, except *θ*_*l*_ = *π* for 1 ≤_*l*_ ≤ *L*_1_ and *θ*_*l*_ = 3*π/*2 for *L*_1_ *<* _*l*_ 2*L*. Then, genotype 00 … 0 is the sole genotype that encodes the phenotypic optimum, and any mutation 1 → 0 is beneficial with the fitness effect given by equations (5), (6). In this model, all genotypes with trait value *x*_*i*_ have access to *b*_*i*_ = |*x*_*i*_|*/δ* beneficial mutations that improve trait *i*.

#### 2.1.6 Nested Fisher’s geometric model (nFGM)

In all GPMs described above, we assumed that both traits *x*_*i*_ are additive and all mutations have exactly the same phenotypic effect *δ*. Here, we consider a more complex “nested” Fisher’s geometric model (nFGM), in which each “high-level” functional trait *x*_*i*_ is decomposed into *n*_*i*_ “low-level” functional traits 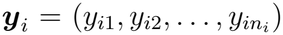, with 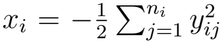 (Figure 5A). We assume that the lower-level traits *y*_*ij*_ are additive and tightly integrated, such that every site on chromosome *i* contributes to every *y*_*ij*_ via equations that generalize equation (2) to *n*_*i*_ dimensions, 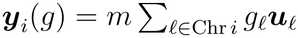, where *m >* 0 is the contribution of each site and ***u***_*l*_ are distributed uniformly on the unit sphere in 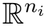 . When the length *L*_*i*_ of chromosome *i* is large, the central limit theorem dictates that, across random genotypes and random directions ***u***_*l*_, the values *y*_*ij*_ are uncorrelated and approximately normally distributed with zero mean and variance *σ*_*i*_ = *L*_*i*_*m*^2^*/*(2*n*_*i*_). Thus, the majority of genotypes are located within the sphere of radius 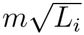 of the optimal phenotype ***y***_*i*_ = **0**, i.e., this optimal phenotype is located in the genotype-dense region of the space of low-level traits, similar to the pleiotropic genotype-phenotype map with a high-entropy optimum described above. However, since *x*_*i*_ is a quadratic function of ***y***_*i*_, −2*x*_*i*_*/σ*_*i*_ is distributed as *χ*^2^ with *n*_*i*_ degrees of freedom, which in turn implies that, for *n*_*i*_ ≥ 3, the peak of the distribution of the high-level trait *x*_*i*_ among genotypes does not coincide with the optimal value *x*_*i*_ = 0. This in turn suggests that the fraction of mutations that improve performance of the high-level module *i* should decline for genotypes with smaller |*x*_*i*_|, and this is indeed the case (see below). Thus, the nFGM model is more similar to the modular GPFM with a low-entropy optimum than the one with a high-entropy optimum.

In the vicinity of radius 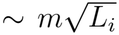 of the optimal phenotype ***y***_*i*_ = **0**, each mutation affecting trait *x*_*i*_ can be approximated by a random vector ***m***_*i*_ with magnitude ∥***m***_*i*_∥ = *m* and a uniformly distributed direction. Thus, if a mutation with effect ***m***_*i*_ occurs in an organism with the trait vector ***y***_*i*_, the mutant’s trait vector is ***y***_*i*_ + ***m***_*i*_, as in the canonical FGM (Tenaillon, 2014). Denoting the resulting effect of this mutation on the functional trait *x*_*i*_ by _Δ*i*_, it can be shown that _Δ*i*_ is approximately normally distributed with mean −*m*^2^*/*2 and standard deviation 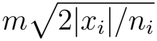 (Tenaillon, 2014). To make the nested FGM directly comparable to the modular GPFM with a low-entropy optimum, we set 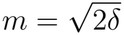 so that the mean effect of a mutation on *x*_*i*_ is −*m*^2^*/*2 = −*δ*, matching the fixed effect size in the modular GPFM. With this calibration, the distribution of mutational effects on *x*_*i*_ has mean −*δ* and standard deviation 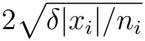 (Figure S4). Consequently, the fraction of module-improving mutations declines with decreasing |*x*_*i*_|, as expected for a model with a low-entropy optimum. Importantly, and in contrast to the modular GPFM with a low-entropy optimum, this decline is non-linear (Figure 5B). Furthermore, in contrast to all models discussed above, the magnitude of trait improvements also declines non-linearly with declining |*x*_*i*_| (Figure 5C). Therefore, we can think of nFGM as probing the robustness of our results with respect to the assumption of trait additivity and the assumption that all mutations have exactly the same phenotypic effect *δ*.

### 2.2 Models of evolutionary dynamics

The key population-genetic parameters that determine the evolutionary dynamics on our GPFMs are the mutation rate *µ* (per site per generation), the reassortment rate *ρ* (per individual per generation) and the population size *N* . We denote the total genome-wide mutation rate by *U* = 2*µL* (per generation), the genome-wide beneficial mutation rate by *U*_*b*_, and the rate of beneficial mutations on chromosome *i* = 1, 2, by *U*_*bi*_. For simplicity, we only consider reassortment between chromosomes and ignore recombination within chromosomes. The list of parameters and their values used in simulations are provided in Table 1.

We consider evolutionary dynamics in three regimes. (1) The *successive mutations* regime (also known as the “strong-selection weak-mutation”, or SSWM, regime) is defined by *NU*_*b*_ ≪ 1, such that at any moment in time, at most one beneficial mutation is segregating in the population (Gillespie, 1991; Desai and Fisher, 2007; Kryazhimskiy et al., 2009; Good and Desai, 2015). In this regime, reassortment plays no role. The *concurrent mutations* regime is defined by *NU*_*b*_ *>* 1, such that multiple beneficial mutations segregate in the population at any moment (Desai and Fisher, 2007). In the concurrent mutations regime, we consider two extreme cases with respect to reassortment. Regime (2) is when reassortment is absent (*ρ* = 0) and regime (3) is when chromosomes reassort at every reproduction event (*ρ* = 1).

The focus of our investigation is on the evolutionary trajectories in the trait space for an average adapting population. These average trajectories are described by the equation

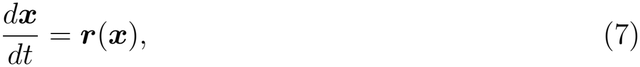

where ***r***(***x***) = *r*_1_(***x***), *r*_2_(***x***) is the vector of instantaneous rates of trait change at the trait-space location ***x***. In particular, we are interested in how the balance between the two module performances changes during a bout of adaptation in a constant environment. We quantify this balance by the module performance ratio *R*(*t*) = *x*_2_(*t*)*/x*_1_(*t*), which measures the relative proximity of the two modules to their respective optima, such that *R* = 1 indicates that both modules are equally close to their optima. It follows from equations (7) that *R* changes along the expected evolutionary trajectory according to equation

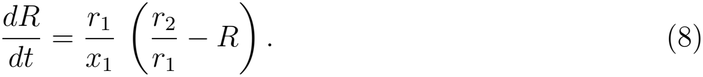

### 2.3 Analytical calculations

We obtain analytical approximations for the average trajectories in the trait space in the successive mutations for the pleiotropic GPFM with a high-entropy optimum and for modular GPFMs with either high- or low-entropy optima. To examine how the module-selection balance observed in the modular GPFM with a low-entropy optimum depends on the supply of adaptive mutations and genetic linkage, we also obtain some analytical results for the concurrent mutations regime in asexual and sexual populations.

#### 2.3.1 Pleiotropic GPFM with a high-entropy optimum

When the population is sufficiently far from the optimum, i.e., *δ* ≪ |*x*_*i*_ |, the selection coefficients of mutations are given by expression (4), i.e., the mutation with effect vector ***δ*** is beneficial whenever its projection on the fitness gradient is positive or, equivalently, whenever *θ* (***x***) *< θ < θ* (***x***) + *π*, where 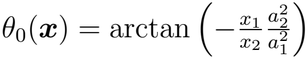 .

For the pleiotropic GPFM with a high-entropy optimum, we carry out analytical calculations in the successive mutations regime, and use it as a comparative reference in other regimes. Mutations with an angle in the infinitesimal interval (*θ, θ* + *dθ*) arise at rate *NUdθ/*2*π*. Such a mutation has selection coefficient *s*(***x***, *θ*) given by equation (4). If the mutation is beneficial, it fixes with probability 2*s*(***x***, *θ*) and it is lost otherwise. A beneficial mutation improves trait 1 by *δ* cos *θ*. Therefore, the expected rate of change in this trait is

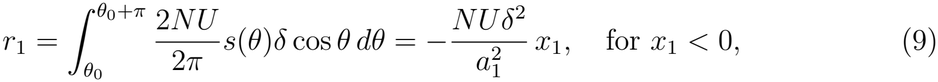

*r*_1_ = and, analogously,

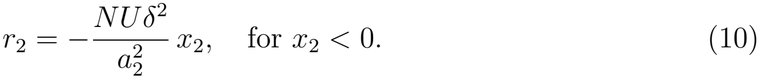

Substituting expressions (9), (10) into equation (7), we obtain

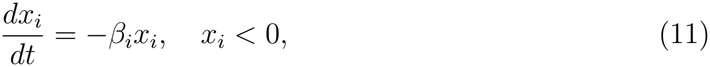

with 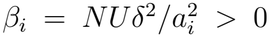 . Equation (11) implies that the population is approaching the optimum along the fitness gradient 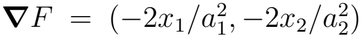 . The solution of equation (11) is given by

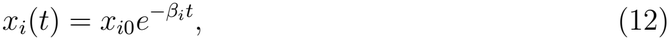

where *x*_*i*0_ is the initial value of trait *i*. Eliminating *t*, we obtain the expression (23) for the expected evolutionary trajectory in the trait space, which implies that the population approaches the optimum along the trait axis that corresponds to the module under weaker selection.

#### 2.3.2 Modular GPFM with a high-entropy optimum

On a modular GPFM with a high-entropy optimum, the distribution of phenotypes among genotypes is approximately normal with mean 0 and variance *Lδ/*2. For genotypes with *x*_1_ *<* 0, the number of mutations that improve trait 1 equals the number of sites with angle *θ*_*l*_ = *π* (those pointing away from the optimum) that are in state 1, plus the number of sites with angle *θ*_*l*_ = 0 (those pointing towards the optimum) that are in state 0, and analogously for trait 2. Then, the number of trait-improving mutations available to a genotype with trait value *x*_*i*_ *<* 0 is approximately normal with mean ⟨*b*_*i*_⟩ = *L/*2 + |*x*_*i*_|*/*2*δ* and variance *L/*8.

We consider the regime 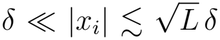 . In this regime, trait-increasing mutations in module *i* arise at rate approximately *NµL/*2 per generation (to the leading order) and have selection coefficient *s*_*i*_(***x***) given by equation (6). In the successive mutations regime, the fraction 2*s*_*i*_ of them go to fixation, and each fixed mutation improves the respective trait by *δ*. Therefore, the expected rate of improvement of module *i* is *r*_*i*_ ≈ *NµLs*_*i*_*δ*. Using expression (6) for *s*_*i*_ in the regime *δ*≪| *x*_*i*_| and the fact that *U* = 2*µL* is the total mutation rate, we find that a typical population in the successive mutations regime moves along a trajectory defined by equation (11). Thus, on a modular GPFM with a high-entropy optimum, an average population also approaches the optimum approximately along the fitness gradient, and its adaptive trajectory in the trait space is approximately given by expression (23).

#### 2.3.3 Modular GPFM with a low-entropy optimum

##### Successive mutations regime

On a modular GPFM with a low-entropy optimum, beneficial mutations in module *i* arise at rate *Nµb*_*i*_ = *Nµ*|*x*_*i*_|*/δ* per generation and have selection coefficient *s*_*i*_(***x***) given by equation (6). In the successive mutations regime, the fraction 2*s*_*i*_ of them go to fixation, and each fixed mutation improves the respective trait by *δ*. Therefore, the expected rate of improvement of module *i* is *r*_*i*_ = 2*Nµ* |*x*_*i*_| *s*_*i*_. Using expression (6) for *s*_*i*_ in the regime *δ* ≪ |*x*_*i*_|, we obtain

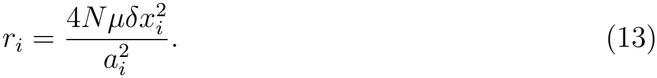

Substituting expression (13) into equation (7), we find that a typical population in the successive mutations regime moves along a trajectory defined by equations

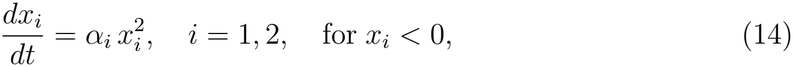

where 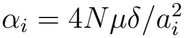 . After eliminating *t*, we obtain

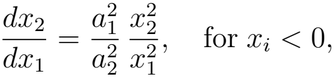

whose solution is given by equation (25) discussed in the Results section. Equation (14) can also be solved directly to yield

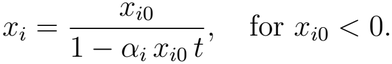

##### Concurrent mutation regime with complete linkage

Here, we show that the line of equal fitness benefits *L*^*\**^ defined by equation (28) is a fixed point of equation (8) for the dynamics of the module performance ratio *R* in the concurrent mutations regime. To demonstrate this, we first denote all quantities evaluated on line *L*^*\**^ with an asterisk. On the line of equal fitness benefits, beneficial mutations arise at rate 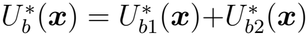, where 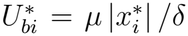 is the rate of adaptive mutations in module *i* along *L*^*\**^, and each provides a fitness benefit *s*^*\**^(***x***) relative to the parent. Then, assuming that the population is at the beneficial mutation-selection quasi-equilibrium, its mean fitness increases at rate 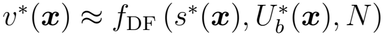, with

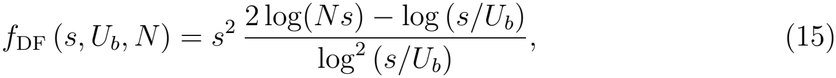

which holds when *s* ≫ *U*_*b*_ (Desai and Fisher, 2007). Gomez et al. (2020) showed that, if the fitness benefits of mutations in two modules are identical (as is the case on *L*^*\**^), the rate of fitness gains 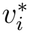 attributable to mutations in module *i* is proportional to 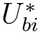 . More precisely, 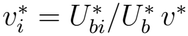, which implies that the rate of change in module performance is

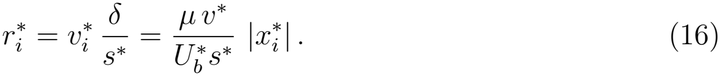

Therefore, 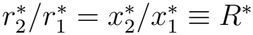, which, according to equation (8), implies that *dR/dt* = 0 at any point on *L*^*\**^. Thus, the line of equal fitness benefits defined by equation (28) is indeed a fixed point for the dynamics of the module performance ratio *R*.

To characterize how populations approach the module-selection balance line in the concurrent mutations regime with complete linkage, we define *v*_*i*_, *i* = 1, 2 to be the rate of fitness gains that is attributed to mutations improving trait *i*. We assume that the population is at the local beneficial mutation-selection balance (Desai and Fisher, 2007), which is determined by the current supplies of beneficial mutations in both modules *U*_*bi*_(***x***) = *µb*_*i*_(***x***) = *µ* |*x*_*i*_| */δ*, their selection coefficients *s*_*i*_(***x***), which are given by equation (6), and the population size *N* .

We approximate *v*_*i*_ (and hence *r*_*i*_ = *v*_*i*_*δ/s*_*i*_) based on the results of Gomez et al. (2020). To this end, we first define 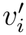 to be the rate of adaptation of module *i* “in isolation”, i.e., the rate of adaptation of an organism with a single trait *i*. As mentioned above, Desai and Fisher (2007) have shown that at the beneficial mutation-selection balance, 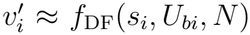 . In the modular GPFM with a low-entropy optimum, *U*_*bi*_ decreases in proportion to |*x*|_*i*_, while *s*_*i*_ does so approximately when |*x*_*i*_| ≫ *δ*, so their ratio remains nearly constant away from the optimum. Gomez et al. (2020) demonstrated that, if module *j* supports a much faster rate of fitness improvements in isolation than module *i* (mathematically, if 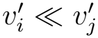 ), then the rate of adaptation of module *j* in the two-module organism is approximately the same as its rate in isolation 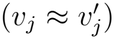, whereas the slower module stalls 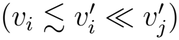 . Moreover, they showed that the rates of trait evolution in a two-trait organism ***v*** = (*v*_1_, *v*_2_) depend on the rates of evolution of the two isolated traits 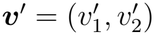 and are nearly independent of the specific combination of parameters *U*_*bi*_ and *s*_*i*_, *i* = 1, 2, that yield a particular ***v***^*′*^. We confirmed these findings using simulations (see Section 2.1 in Supplementary Information).

Given these results, we approximate the rates of improvement of trait *i* as

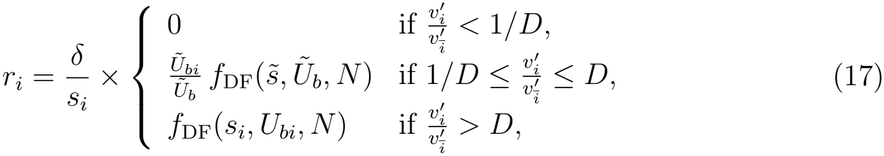

where 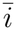 denotes the module other than *i*, 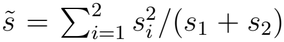 and 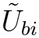 is the effective beneficial mutation rate defined as a solution of equation

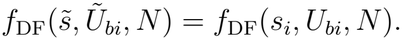

Here 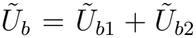 is the total effective beneficial mutation rate. *D* denotes the ratio threshold, which we set to be *D* = 100 although prediction quality is largely insensitive to this choice as long as *D* ≫ 1 (Figure S3). The motivation for the approximation (17) as well as its validation are provided in Section 2.2 in Supplementary Information.

#### Concurrent mutation regime with free reassortment

We model evolution in this regime by assuming that the two modules evolve independently. In this case, since *U*_*bi*_ = *µ* |*x*_*i*_| */δ* and *s*_*i*_ is given by equations (6), we have 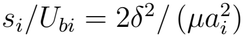, which implies

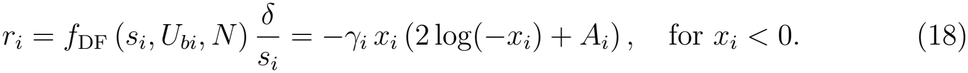

where *f*_DF_ is given by equation (15), and

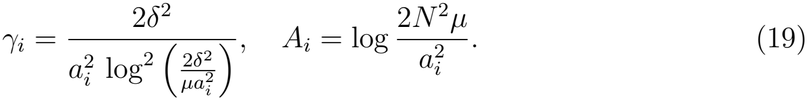

Substituting expression (18) into equation (7), we find that

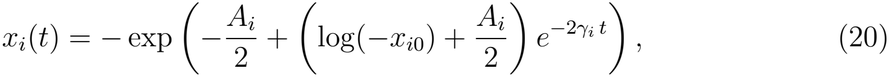

which holds as long as each module adapts in the concurrent mutation regime, i.e., as long as the population stays sufficiently far from the phenotypic optimum, such that 2 log(−*x*_*i*_) + *A*_*i*_ *>* 0. Then, the expected trajectory in the trait space is given by equation

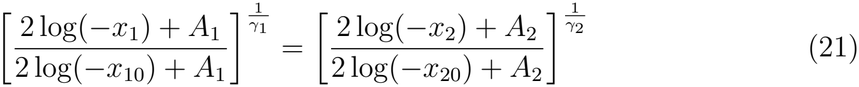

and the module performance ratio is given by

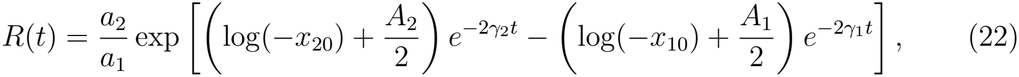

which implies that *R* eventually converges to *a*_2_*/a*_1_. However, since *γ*_*i*_ ∼ *δ*^2^, this convergence is slow (compare to equation (14) where *α*_*i*_ *δ*), and populations may leave the concurrent mutations regime where this approximation is valid before they approach this equilibrium.

### 2.4 Simulations

We carried out two types of evolutionary simulations. For simulations in the successive mutations regime, we assumed that the population is monomorphic and modeled its genotype (or, in the modular GPFMs and the nFGM, its trait values) as a Markov chain using the Gillespie algorithm (Gillespie, 1977). In the concurrent mutations regime, with or without reassortment, we simulated evolution using the Wright-Fisher model (Ewens, 2004). Unless otherwise noted, parameter values used in our simulations are provided in Table 1.

#### 2.4.1 Numerical implementation of different GPFMs

The pleiotropic GPFMs, modular GPFMs, and the nFGM differ in whether explicit geno-types are tracked and in how mutations are implemented, and these differences shape the simulation approach for each model.

##### Pleiotropic GPFMs

In the pleiotropic GPFMs with either low- or high-entropy optimum, genotypes are tracked explicitly, such that each individual in the population is characterized by an explicit binary genome of 2*L* = 400 loci. In each simulation, each locus _*l*_ is assigned a fixed pleiotropic angle *θ*_*l*_, drawn once prior to simulation from Uniform[0, 2*π*) (high-entropy optimum) or Uniform[*π*, 3*π/*2] (low-entropy optimum) and shared across all replicates and evolutionary regimes. Mutations at each locus arise at rate *µ* per individual per generation. A mutation at locus _*l*_ flips its allele (0 ↔ 1) and correspondingly displaces the phenotype by ±***δ***_*l*_ as described in Section 2.1.2.

##### Modular GPFMs

In both modular GPFMs, we do not track genotypes explicitly, but only the trait values *x*_*i*_. In the modular GPFM with the low-entropy optimum, trait value *x*_*i*_ contains all the information needed to carry out the simulations. Indeed, since the phenotypic effects at all loci are the same in our model, we only need to know the number *b*_*i*_ of loci that carry allele 1 and therefore can supply a mutation that increases *x*_*i*_, but which specific loci carry allele 1 is irrelevant. In this model, *b*_*i*_ = |*x*_*i*_|*/δ*, meaning that all the relevant information is indeed contained in *x*_*i*_.

In the GPFM with a high-entropy optimum, the trait value *x*_*i*_ does not fully determine the supply of trait-increasing mutations. As discussed in Section 2.3.2, the number of trait-increasing mutations available to a genotype with trait value *x*_*i*_ is approximately normally distributed with mean ⟨*b*_*i*_ ⟩= *L/*2 − *x*_*i*_*/*2*δ* and variance *L/*8. Moreover, since *L* ≫ 1, the variation around the mean can be ignored without causing much error. Thus, to simplify our simulations further, we assume that every genotype with trait *x*_*i*_ has access to exactly *b*_*i*_ = *L/*2 − *x*_*i*_*/*2*δ* trait-increasing mutations.

In both GPFMs, mutations that increase *x*_*i*_ arise at rate *µb*_*i*_ per individual and shift the trait value *x*_*i*_ by +*δ*; mutations that decrease it arise at rate *µ* (*L*_*i*_ − *b*_*i*_) and shift *x*_*i*_ by −*δ*.

##### Nested FGM

In the nested FGM, mutations in module *i* arise at rate *U/*2, and each such mutation shifts the trait *x*_*i*_ by _Δ*i*_, drawn from the normal distribution with mean −*m*^2^*/*2 and variance 2|*x*_*i*_|*m*^2^*/n*_*i*_ (see Section 2.1). We set 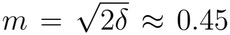 and *n*_*i*_ = 10. Reassortment acts as in the modular GPFM, by swapping the trait *x*_2_ between two individuals in a mating pair.

#### 2.4.2 Simulations in the successive mutations regime

In this regime, the population is assumed to remain monomorphic between fixation events, with transients to fixation treated as instantaneous. We assume that neutral or deleterious mutations never fix. Therefore, we draw the waiting time *τ* until the appearance of the next beneficial mutation from an exponential distribution with rate *NU*_*b*_, where *U*_*b*_ is the genome-wide beneficial mutation rate, which is model-dependent (see below). We then determine the selection coefficient *s* of the arisen beneficial mutation, which is also model-dependent, and evaluate its probability of fixation using Kimura’s expression *P*_fix_(*s*) = (1 − *e*^*−*2*s*^)*/*(1 − *e*^*−*2*Ns*^) (Gillespie, 2004). If the mutation is lost, time advances by *τ* but the state of the population is unchanged; if it fixes, the state of the population is updated.

##### Pleiotropic GPFMs

In pleiotropic GPFMs, we determine *U*_*b*_ as follows. We first evaluate, for every locus _*l*_, the selection coefficient *s*_*l*_ of a mutation that flips its allele, using expression (3) with the displacement +***δ***_*l*_ or −***δ***_*l*_ determined by the allele currently at locus _*l*_. We form the set of beneficial loci *B* = {_*l*_ : *s*_*l*_ *>* 0 } as those with positive selection coefficients. The number of available beneficial mutations *b* is the size of this set, *b* = |*B*|, and *U*_*b*_ = *µb* per individual per generation. Once a beneficial mutation arises, we draw a random locus _*l*_ from *B* which yields the selection coefficient *s*_*l*_ of the arisen mutation.

##### Modular GPFMs

Here, *U*_*b*_ = *µ* (*b*_1_ + *b*_2_). We draw the chromosome *i* on which the beneficial mutation occurs with probability proportional to *b*_*i*_. The mutant’s trait *x*_*i*_ is then increased by *δ*, and its selection coefficient *s*_*i*_ is evaluated according to equation (5).

##### nFGM

We have *U*_*b*_ = *U* (*P*_1_ + *P*_2_)*/*2, where *P*_*i*_ = Pr(_Δ*i*_ *>* 0) is the probability that a mutation in module *i* improves trait *i*, computed using the normal distribution with mean −*δ* and standard deviation 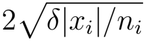 . The module *i* in which the beneficial mutation occurs is chosen with probability proportional to *P*_*i*_. Then, the phenotypic effect _Δ*i*_ is drawn from the same normal distribution, conditional on _Δ*i*_ *>* 0. The selection coefficient is calculated from the resulting phenotype, and proposals with non-positive selection coefficients are rejected. Note that in the exact nFGM, 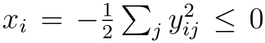, but because we approximate _Δ*i*_ by a normal distribution, a mutation can occasionally overshoot the optimum and make *x*_*i*_ positive. In this case, we assume that the module has effectively reached the optimum and *x*_*i*_ stays at this small positive value for the rest of the simulation. Adaptation may continue in the other module until the mean fitness reaches the termination condition. Nevertheless, such cases are very rare (≈0.2% of all recorded states in these simulations), and the resulting positive values are small (median 0.01, at most ≈ *δ*).

#### 2.4.3 Wright-Fisher model simulations

In the Wright-Fisher model simulations, each generation contains three stages: mutation, reassortment, and selection/reproduction. Implementation of mutation and reassortment varies across GPFM models and is discussed below. After mutations and reassortment events have occurred and the traits and fitness *F* of all individuals have been calculated, the selection/reproduction step is applied uniformly across all models by sampling the number of descendants of each individual from a multinomial distribution with *N* trials and probabilities proportional to individual Wrightian fitness *W* = *e*^*F*^ .

##### Pleiotropic GPFMs

In both pleiotropic GPFMs, the number of mutation events per generation is drawn from the Poisson distribution with mean *NU* . For each event, an individual who received the mutation and the locus at which the mutation occurs are drawn uniformly. Then, the allele at the specified locus is flipped and the phenotypes and fitness are updated accordingly. For *ρ* = 1, all individuals are grouped into pairs, with every possible pairing equally likely, and exchange their second chromosome containing loci *L*_1_ + 1, …, 2*L*.

##### Modular GPFMs

Here, each individual *j* acquires mutations that increase *x*_*i*_ at rate *µb*_*ij*_ and mutations that decrease *x*_*i*_ at rate *µ*(*L*_*i*_ − *b*_*ij*_), where *b*_*ij*_ is the value of *b*_*i*_ for individual *j* (Section 2.4.1). We implement this in two steps. First, we draw the total numbers of mutations on chromosome *i* that increase and decrease *x*_*i*_ in the population from the Poisson distributions with means 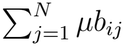 and 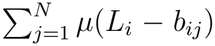, respectively. Second, we assign each mutation that increases *x*_*i*_ to an individual drawn, without replacement, with probability proportional to its *b*_*ij*_, and shift its *x*_*ij*_ by +*δ*; mutations that decrease *x*_*i*_ are assigned in the same way, with probabilities proportional to *L*_*i*_ − *b*_*ij*_, and shift *x*_*ij*_ by −*δ*. For *ρ* = 1, all individuals are grouped into pairs, with every possible pairing equally likely, and the trait values *x*_2_ are swapped within each pair.

##### nFGM

In the nested FGM, the number of mutations per module per generation is drawn from the Poisson distribution with mean *NU/*2. The individuals that receive these mutations are drawn uniformly without replacement, and each mutation changes the trait by _Δ*i*_, drawn from the normal distribution with mean −*δ* and standard deviation 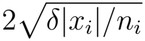 . Because this distribution is an approximation, a mutation can occasionally make *x*_*i*_ positive, which is impossible in the exact nFGM. In contrast to the successive mutations regime, where such an overshoot can occur at most once per module, here overshoots can occur in any generation and be inherited, so we set positive values to zero to keep them from building up in the population. For *ρ* = 1, all individuals are grouped into pairs, with every possible pairing equally likely, and *x*_2_ is swapped within each pair. However, it can be shown that the probability that a mutation overshoots the optimum, Pr(_Δ*i*_ *>* |*x*_*i*_| ), is at most 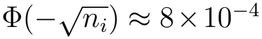 for *n*_*i*_ = 10, where Φ is the standard normal cumulative distribution function.

#### 2.4.4 Initialization and termination

Unless otherwise noted, all simulations are initialized at six locations in the trait space, with target fitness *F*_0_ = log(0.25) ≈ −1.39 and different target module performance ratios *R*_0_ = ∈ {*x*_20_*/x*_10_ 0.16, 0.31, 0.625, 1.25, 2.5, 5} as described below for different models. Simulations are terminated when the population’s mean Wrightian fitness reaches 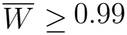 . We ran 250 replicate simulations per initial condition.

##### Pleiotropic GPFMs

Genotypes with nearly the same phenotype can differ in which mutations are available to them, so the choice of the starting genotype can affect the subsequent dynamics. To avoid biasing this choice, we draw the starting genotype from the uniform distribution over the set A_*ϵ*_(***x***_0_) of all genotypes whose phenotypes lie within Euclidean distance *ϵ* = 0.02 of the target phenotype ***x***_0_. We sample genotypes from this set using the maximum-entropy distribution, as follows.

We first generate candidate genotypes whose phenotypes equal ***x***_0_ on average. In each candidate genotypes, allele 1 at locus _*l*_ is independently drawn with probability *p*_*l*_ = [1 + exp ⟨***λ, u***_*l*_⟩]^*−*1^, where ***u***_*l*_ = (cos *θ*_*l*_, sin *θ*_*l*_)^*T*^ is the unit vector at site _*l*_, and the vector ***λ*** = (*λ*_1_, *λ*_2_)^*T*^ is found by numerically solving *δ* ∑ _*l*_ *p*_*l*_***u***_*l*_ = ***x***_0_. Among all probability distributions over genotypes whose mean phenotype is ***x***_0_, this one has the maximum entropy (see Section 3 in the Supplementary Information). We generate candidates in batches of 5,000 and keep those whose phenotypes fall within distance *ϵ* of ***x***_0_, until at least 200 candidates have been kept or 4 × 10^6^ candidates have been generated (this limit was never reached). We randomly draw one of these candidates with probability proportional to exp [⟨***λ, x***(*g*) −***x***_0_ ⟩*/δ*], where ***x***(*g*) is the trait vector of candidate genotype *g*. The selected initial genotype is then used to initialize all replicate populations for that initial condition in all three evolutionary regimes.

##### Modular GPFMs

In both modular GPFMs, the initial population is monomorphic with phenotype values *x*_*i*0_ obtained as the nearest *δ*-multiple and clamped to *x*_*i*0_ ≤ 0.

##### nFGM

Here, the initial population is monomorphic at ***x***_0_.

#### 2.4.5 Visualization of evolutionary trajectories

##### Trait-space trajectories

To visualize variation among replicate trajectories in the trait space, we measure how far replicates deviate from the mean trajectory in the direction perpendicular to it. Since replicates take different numbers of generations to reach the termination criterion, we compare them at the same fraction of their own trajectories. Specifically, in each simulation, we record a sequence of population-mean trait vectors, one per generation in the Wright-Fisher simulations and one per fixation event in the successive-mutations simulations. For each replicate, we sample 20 trait vectors from this sequence, uniformly spaced from first to last. The mean trajectory is obtained by averaging the first, second, …, 20th sampled trait vectors across replicates and connecting these 20 points.

We visualize the variation around the trajectory as follows. First, at each of the 20 sampled points along the mean trajectory, we approximate its tangent by the line through the two neighboring mean points or the adjacent point at either end. Then, we project each replicate’s deviation from the mean onto the orthogonal direction. We then estimate the standard deviation of these projections across replicates. The shaded bands in the trait-space panels of Figures 2–4 show the mean trajectory plus or minus one standard deviation in this perpendicular direction.

**Figure 2.**
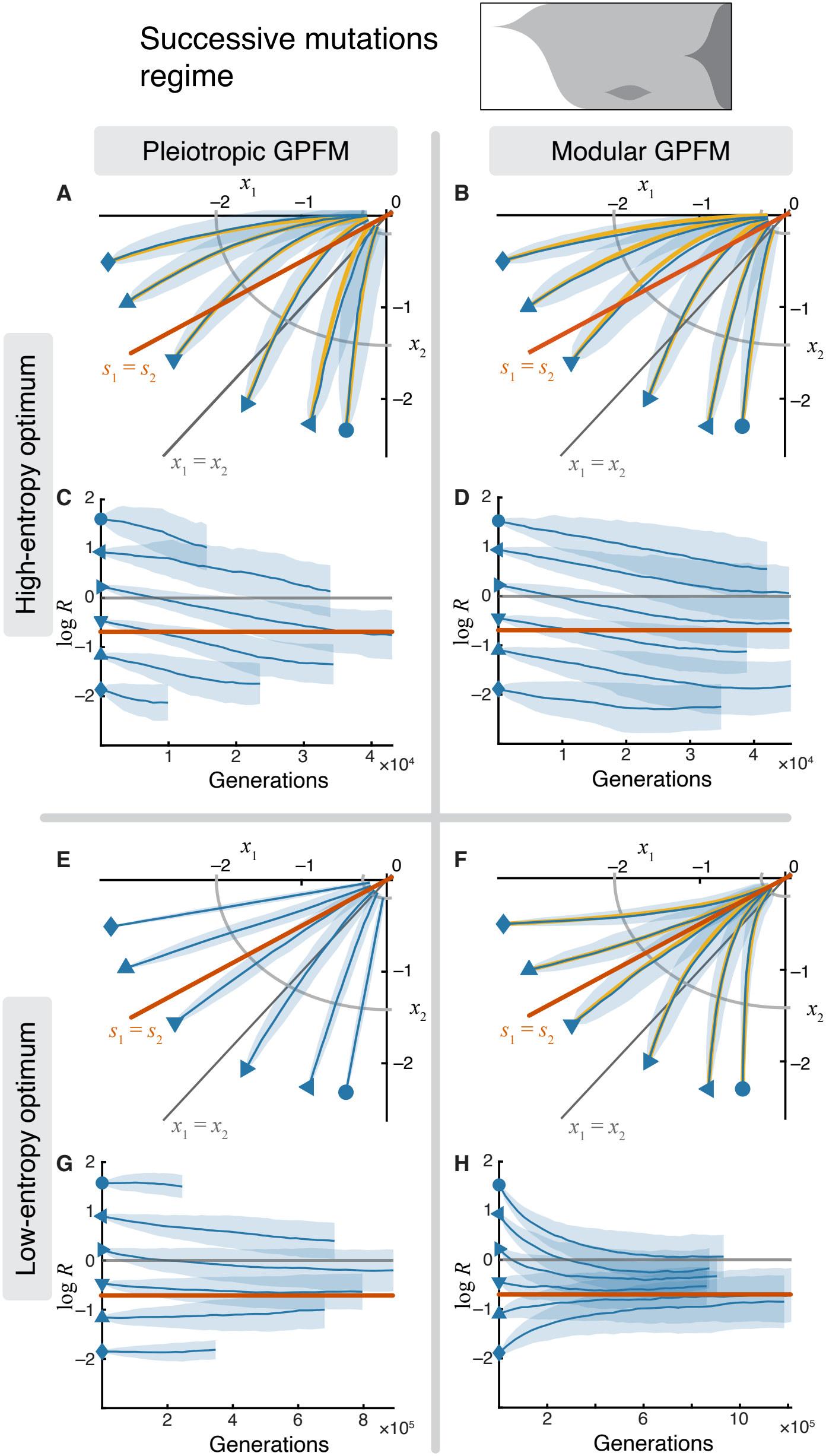
Trait evolution in the successive mutations regime (previous page). A,B,E,F. Evolutionary trajectories in the trait space on different GPFMs, as indicated. Populations are initialized at different locations in the trait space with initial fitness near *F*_0_ = −1.39 (see Section 2.4.4 for details). Yellow lines show analytical predictions where available (equation (23) for panels A,B; equation (25) for panel F). Blue lines show simulated trajectories averaged over 250 replicates; shaded bands show ±1 standard deviation across replicates computed as described in Section 2.4. Gray line is the diagonal *x*_1_ = *x*_2_. Orange line is the line of equal fitness benefits (equation (26)). **C, D, G, H**. Module performance ratio *R* = *x*_2_*/x*_1_ over time for the respective GPFMs. Solid lines show the mean and shaded bands show ±1 standard deviation of log *R* across replicates (see Section 2.4 for details). Each curve ends when more than 20% of replicates have a trait within *δ* of its optimum. Orange horizontal line indicates the module performance ratio 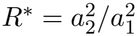 predicted at the module-selection balance. The simulation parameters are *N* = 10^4^, *δ* = 0.1, 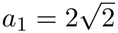, *a*_2_ = 2, *L*_1_ = *L*_2_ = 200, and *µ* = 5 × 10 per locus per generation (genome-wide *U* = 2 × 10^*−*6^).

**Figure 3.**
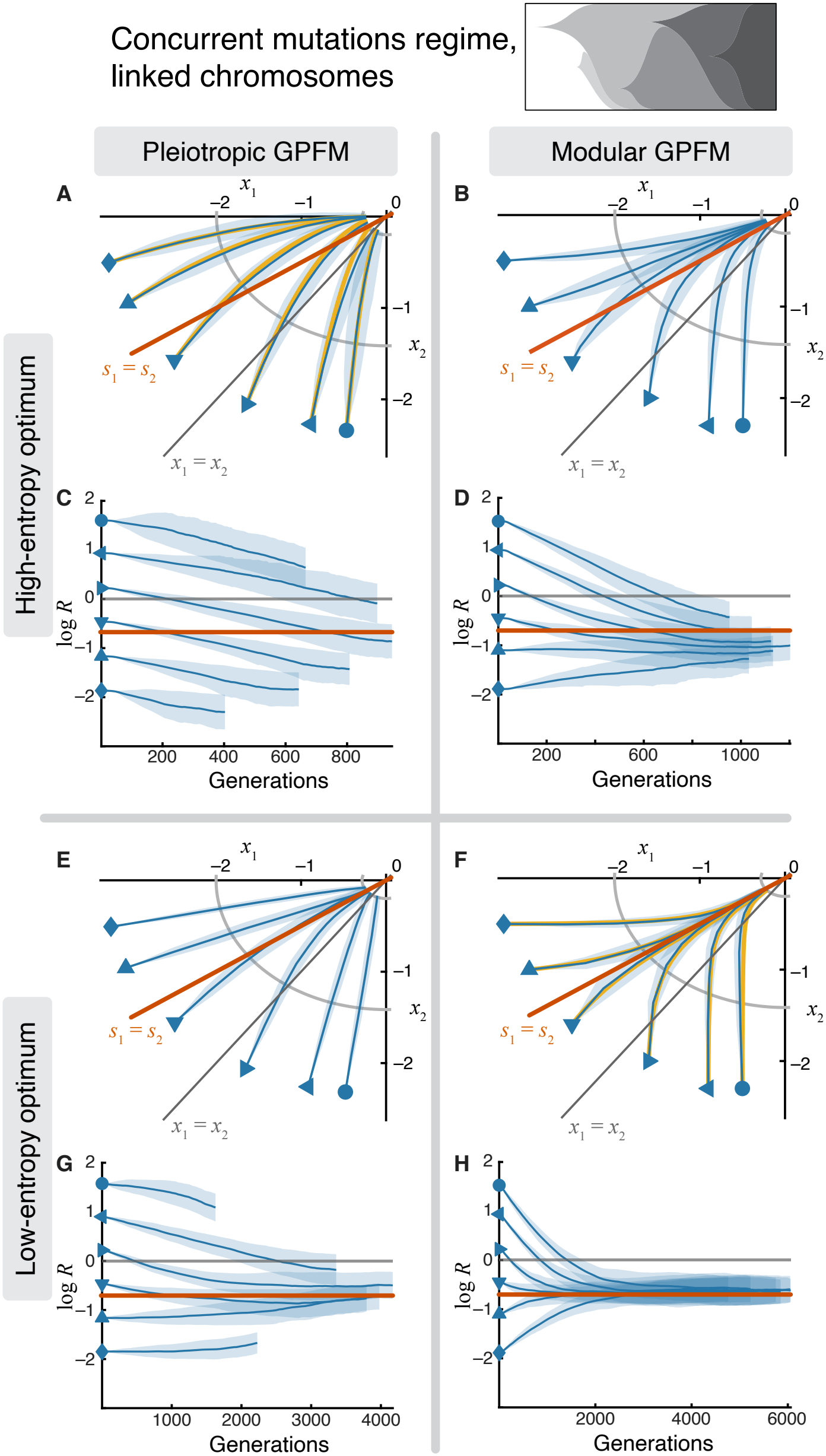
Trait evolution in the concurrent mutations regime with completely linked chromosomes (previous page). A,B,E,F. Evolutionary trajectories in the trait space on different GPFMs, as indicated. Yellow lines show analytical predictions where available (equation (23) in panel A; equation (17) substituted into equation (7) and solved numerically in panel F). **C, D, G, H**. Module performance ratio *R* = *x*_2_*/x*_1_ over time for the respective GPFMs. Notations are the same as in Figure 2 unless otherwise noted. Simulation parameters are the same as in Figure 2, except *µ* = 10^*−*5^ per locus per generation (*U* = 4 × 10^*−*3^) and *ρ* = 0 (no reassortment).

**Figure 4.**
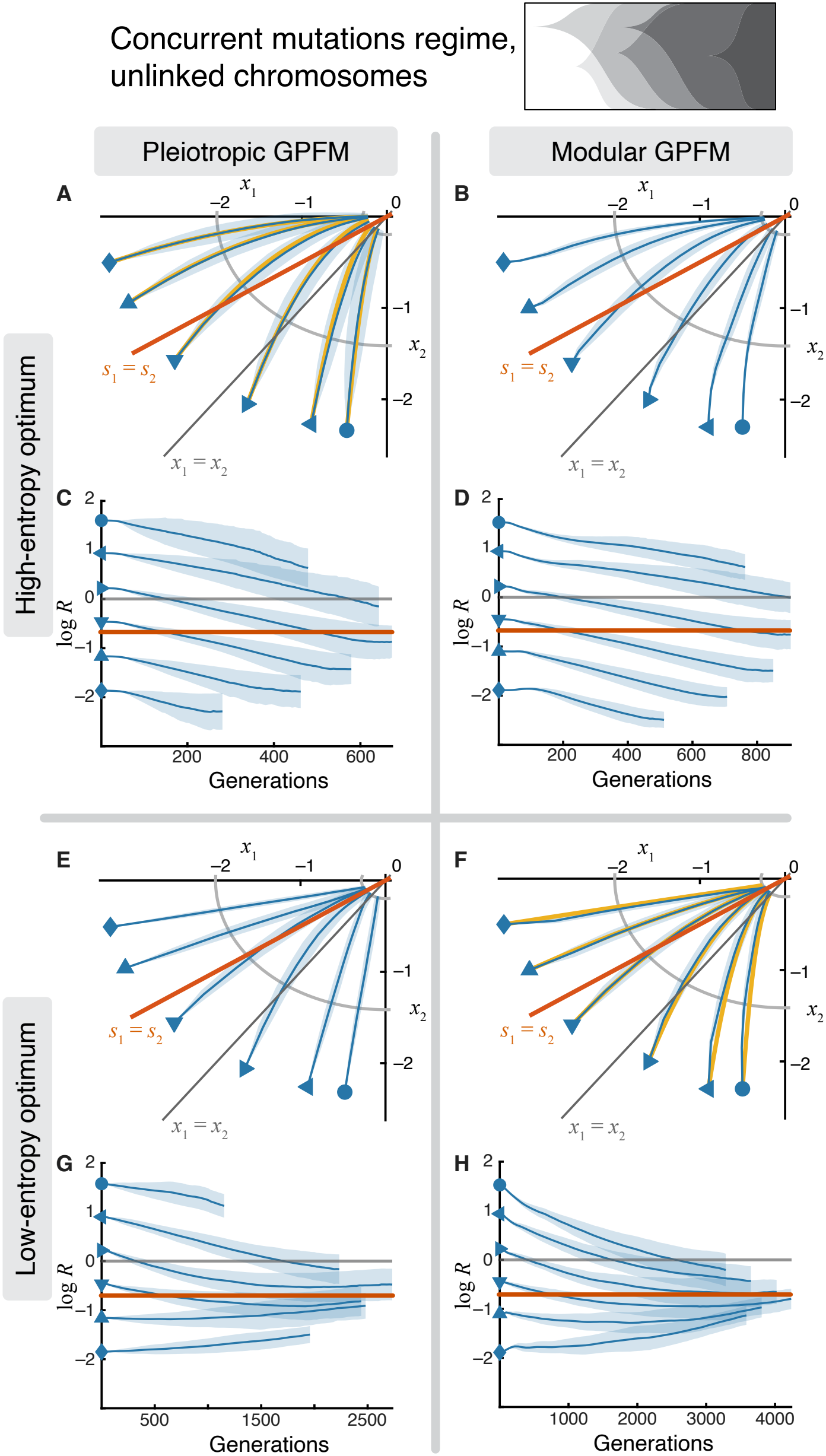
Trait evolution in the concurrent mutations regime with freely reassorting chromosomes (previous page). A,B,E,F. Evolutionary trajectories in the trait space on different GPFMs, as indicated. Yellow lines show analytical predictions where available (equation (23) in panel A; equation (20) in panel F). **C, D, G, H**. Module performance ratio *R* = *x*_2_*/x*_1_ over time for the respective GPFMs. Notations are the same as in Figure 2 unless otherwise noted. Simulation parameters are the same as in Figure 3, except *ρ* = 1 (free reassortment).

##### Module performance ratio

We visualize the natural logarithm of the performance ratio *R* rather than *R* itself because we want to show that *R* declines in our simulations on the pleiotropic GPFM, even when *R*_0_ is already small (e.g., for *R*_0_ = 0.16, and *R*_0_ = 0.31), and these declines are difficult to see on the linear scale. However, in models with high-entropy optima, *x*_*i*_ can switch sign, in which case *R* stops being meaningful. Thus, to avoid this problem, once a replicate reaches a point where min_*i*_ |*x*_*i*_| *< δ*, we exclude it from the calculation of log *R*. However, this procedure restricts the ensemble of populations that contribute to the estimate of mean log *R* and biases this estimate. In particular, since we chose *a*_2_ *< a*_1_ for all our GPFMs (see Table 1), our populations tend to approach the optimal value of trait 2 faster than the optimal value of trait 1, which leads us to preferentially exclude populations with smaller values of *R* from our sample, which biases the estimate of log *R* upwards at later times. Furthermore, since different simulations terminate at different times, estimates of mean log *R* become progressively noisier as time goes on. To avoid excessive noise, we display mean log *R* only at time points where at least 80% of the replicates (200 of 250) still contribute.

### 2.5 Analysis of Lenski’s LTEE metagenomic data

We analyzed published metagenomic data from Lenski’s Long-Term Evolution Experiment (LTEE) obtained by Good et al. (2017) and available at https://github.com/benjaminhgood/LTEE-metagenomic. For each of the six non-mutator populations (Ara+1, Ara+2, Ara+4, Ara+5, Ara −5, Ara −6), we extracted all mutations assigned the “PASS” status in the annotated time course files. We excluded intergenic mutations because they cannot be unambiguously assigned to specific genes. We retained a total of 1,464 genic mutations. We used the provided estimates of appearance times *t*_*a*_ of each mutation, which were determined from a hidden Markov model analysis.

#### Sliding-window analysis

To examine how the diversity of genes targeted by selection changes over the course of the experiment, we computed the effective number of gene targets in overlapping windows of the LTEE time course. Each sliding window spans 10^4^ generations and successive windows are offset by 2,500 generations. Mutations from all six populations are pooled within each window. For each window, we compute Simpson’s index 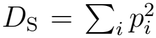, where *p*_*i*_ is the fraction of mutations in gene *i* among all mutations observed in the current window, and report 1*/D*_S_ as the effective number of genes under positive selection. The number of mutations observed in different windows varies between 104 and 326. To control for this unequal variation, we subsampled 104 mutations per window without replacement. We computed *D*_S_ for each subsample and repeated this procedure 2 × 10^3^ times. The plotted effective number of gene targets is the reciprocal of the mean *D*_S_ across subsamples. The lower and upper limits of the shaded bands are the reciprocals of the 97.5th and 2.5th percentiles of *D*_S_, respectively.

#### Multi-hit gene analysis

The PASS filter identifies variants that reached approximately 10% frequency in at least two consecutive sampled time points. As such, this set includes both driver and hitchhiker mutations. Since we do not expect the rate or the genomic distribution of hitchhiker mutations to change over time, we attribute all temporal changes in the distribution of observed mutations to drivers. To assess possible confounding by hitchhikers, we repeated the sliding-window analysis using only multi-hit genes with at least two, three, or four independent mutations. Higher multiplicity thresholds enrich the set for targets of adaptation (Good et al., 2017) but also exclude a larger fraction of genuine targets. The results are shown in Figure S8.

#### Genomic distribution

To visualize how the genomic location of mutations shifts over the course of the experiment, in Figure 6B, we plotted the distribution of genomic positions of all genic PASS mutations separately for an early epoch (*t*_*a*_ ≤ 17,500 generations) and a late epoch (*t*_*a*_ *>* 35,000) generations. The late epoch was chosen as the shortest window that terminates at the final sampling time point and contains at least as many mutations as the early epoch.

### 2.6 Code availability

All the main simulations were performed in MATLAB (Mathworks, Inc.). LTEE analyses were performed in Python. Code is available at https://github.com/mkkim1894/ES_Project.

## 3 Results

To understand how variational modularity and the entropy of the trait space might constrain the evolutionary dynamics at the trait level, we compare these dynamics on four simple genotype-phenotype-fitness maps (GPFMs) that differ in the degree of pleiotropy (highly pleiotropic versus variationally modular) and in how dense the genotype space is in the neighborhood of the optimal phenotypes (high-versus low-entropy optima), as schematically depicted in Figure 1 (see Section 2.1 in “Methods” for details). In all GPFMs, the organism has two functional traits ***x*** = (*x*_1_, *x*_2_), and fitness *F* is a quadratic function of ***x***, as given by equation (1), the genome has 2*L* binary sites, and both traits are additive. We consider mutations that flip the allele at one site at a time, and we assume that the magnitude of the phenotypic effect *δ* of mutations at all sites is the same.

The difference between the pleiotropic and modular GPFMs is that, on the former, all 2*L* binary loci contribute to both traits, whereas on the latter, the first set of *L* loci contribute to functional trait 1 and the second set of *L* loci contribute to functional trait 2 (Figure 1A,B). For simplicity, we refer to these two sets of *L* loci as chromosomes 1 and 2. As a result, in the pleiotropic GPFM, a mutation at site _*l*_ moves an organism by the trait vector ***δ***_*l*_ = ±(*δ* cos *θ*_*l*_, *δ* sin *θ*_*l*_) where *θ*_*l*_ is the trait angle associated with site _*l*_, which is randomly drawn from a uniform distribution (either on [0, 2*π*) in case of a high-entropy optimum (Figure 1C) or on [*π*, 3*π/*2) in case of a low-entropy optimum (Figure 1E); see Sections 2.1.2 and 2.1.3 for details). In other words, in the pleiotropic GPFM, all mutations affect both traits, and different mutations affect the two traits in different proportions. In contrast, in the modular GPFMs, each mutation moves the organism either by trait vector ±***δ***_1_ = (±*δ*, 0) or by trait vector ±***δ***_2_ = (0, ±*δ*). Thus, each mutation affects either trait 1 or trait 2.

The difference between the models with low- and high-entropy optimum is that the phenotypic space in the vicinity of the trait optimum is either sparsely or densely populated by genotypes. In the high-entropy cases (on either pleiotropic of modular GPFMs), the distribution of phenotypic effects of mutations remains nearly constant across the trait space in the neighborhood of the optimum. In contrast, in the low-entropy cases, the distribution of phenotypic effects of mutations varies drastically across the trait space in the neighborhood of the optimum. For example, in the modular GPFM with a high-entropy optimum, all genotypes whose trait values *x*_*i*_ are within roughly 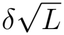 of the optimum have access to approximately 50% of trait-increasing mutations and 50% of trait-decreasing mutations. In contrast, in the modular GPFM with a low-entropy optimum, the fraction of mutations that move a trait value *x*_*i*_ towards the optimum is proportional to *x*_*i*_ (see Section 2.1.5). In other words, when the optimum is low-entropy, organisms exhibit a strong mutational pressure away from it, but they exhibit almost no such pressure when the optimum is high-entropy. Finally, we note that our pleiotropic GFPM with a high-entropy optimum becomes equivalent to the canonical FGM in the limit of a large genome *L* ≫ 1 (see Section 2.1.2).

We compare the evolutionary dynamics that emerge on these four GPFMs in three evolutionary regimes that differ by the total supply of adaptive mutations and the mode of reproduction. In Section 3.1, we begin by considering evolutionary dynamics in the successive mutations regime, also known as the “strong selection weak mutation” regime (Gillespie, 1983; Orr, 2002; Desai and Fisher, 2007; Kryazhimskiy et al., 2009). In this mathematically more tractable limit, adaptive mutations arise so rarely that the population stays mostly monomorphic (deleterious variation is ignored), with occasional instantaneous isolated selective sweeps. Since there is no standing genetic variation, whether the organism reproduces sexually or asexually is irrelevant. If adaptive mutations arise frequently (roughly, at least once per generation), the dynamics transition to the concurrent mutations regime where multiple beneficial alleles segregate simultaneously (Desai and Fisher, 2007). In this regime, which we address in Section 3.2, we consider either an asexual population, in which case both chromosomes remain in complete genetic linkage with each other, or a sexually reproducing one, in which the two chromosomes reassort at every reproduction event. Reassortment within chromosomes is ignored.

### 3.1 Trait dynamics in the successive mutations regime vary qualitatively across GPFMs

#### Simulation results

We begin by simulating the evolutionary dynamics of populations evolving in the successive mutations regime on each of our four GPFMs, starting from different points in the trait space (see Section 2.4 for details and Table 1 for parameter values). We then compute trajectories in the trait space averaged over the simulation ensemble. We find that these trajectories are qualitatively different across our GPFM models (Figure 2). When the fitness optimum is located in a genotype-dense region of the trait space, adaptation starting from different initial conditions on either pleiotropic or modular GPFMs proceeds along trajectories that converge and approach the optimum along the axis that represents the trait under weaker selection (Figure 2A,B). When the fitness optimum is located in a genotype-sparse region of the trait space and the GPFM is highly pleiotropic, adaptive trajectories starting from different initial conditions are close to linear and do not converge other than at the fitness optimum itself (Figure 2E). Finally, on a modular GPFM with a low-entropy optimum, all adaptive trajectories converge and approach the fitness peak along a line that bisects the trait space (Figure 2F).

To further visualize this difference in the dynamics of adaptation, we compute the ratio of functional performances of the two modules *R* = *x*_2_*/x*_1_. On GPFMs with high-entropy optima, this ratio decreases regardless of the initial condition (Figure 2C,D). On the pleiotropic GPFM with a low-entropy optimum, the ratio stays approximately constant (Figure 2G). Finally, on the modular GPFM with a low-entropy optimum, the performance ratio decreases for populations starting at initially high *R* values and increases for populations starting with initially low *R* values (Figure 2H). As a result, the performance ratios converge to an intermediate value on the modular GPFM with a low-entropy optimum, but not on any of the other GPFMs.

#### Analytical approximations

To better understand the qualitative difference in the observed trait dynamics, we obtained analytical approximations for the average evolutionary trajectories for both modular GPFMs and for the pleiotropic GPFM with a high-entropy optimum under the assumption that traits remain sufficiently far from the optimum, |*x*_*i*_| ≫*δ*. Analytical results for the pleiotropic GPFM with a low-entropy optimum could not be easily obtained.

It is straightforward to show that on both GPFMs with a high-entropy optimum (Sections 2.3.1 and 2.3.2 in “Methods“), a typical population approaches the optimum approximately along the fitness gradient-ascent trajectory given by

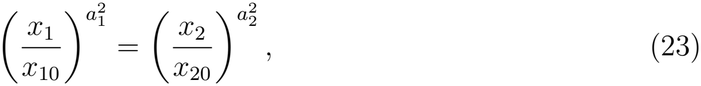

with *a*_*i*_ being the parameter that modulates the strength of selection on trait *i* (with smaller value of *a*_*i*_ indicating stronger selection, see equation (1)) and *x*_*i*0_ *<* 0 being the initial value of trait *i*. It is also easy to show that the module performance ratio *R* = *x*_2_*/x*_1_ changes as

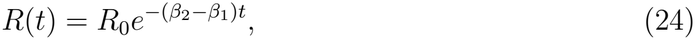

where *R*_0_ = *x*_20_*/x*_10_ is the initial module performance ratio and 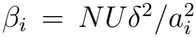, with *N* being the population size and *U* = 2*Lµ* being the genome-wide mutation rate. Equations (23), (24) show that, unless selection pressures on both modules are exactly identical (*a*_1_ = *a*_2_), the module under stronger selection improves at a faster rate. As a consequence, the performance ratio either increases to infinity or declines to zero, and the population approaches the fitness optimum along the trait axis corresponding to the weakly selected module (Figures 2A,B).

Analogous calculations for the modular GPFM with a low-entropy optimum, detailed in Section 2.3.3 in “Methods“, show that a typical population evolves in the trait space along a hyperbolic trajectory,

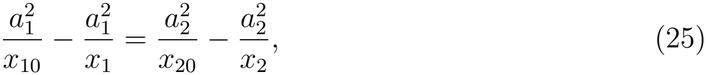

so that all trajectories, irrespective of the initial condition, approach the fitness peak along the line

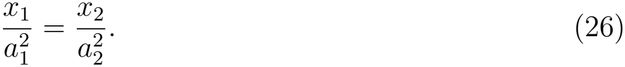

It is also easy to show that, as the organism adapts, the ratio of module performances *R* changes according to equation

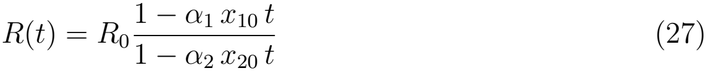

and approaches 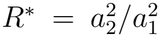 irrespective of the initial condition (Figure 2F,H). Here, 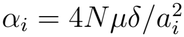 and *µ* is the per-site mutation rate. This implies that, unlike the GPFMs with a high-entropy optimum where the strongly selected functional module always improves faster than the weakly selected one, on the modular GPFM with a low-entropy optimum, both functional modules eventually improve at proportional rates. We refer to the state of the population where both modules improve while their performance ratio remains constant as the “module-selection balance”. At this balance, *r*_2_*/r*_1_ = *x*_2_*/x*_1_ (see equation (8)), meaning that the rates of trait improvement are proportional to the distances to their respective optima.

To better understand this state, consider equation (26) and notice that the fitness effect of a mutation improving module *i* is approximately given by 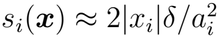 (see equation (6)). Therefore, equation (26), which characterizes the module-selection balance, can be rewritten as

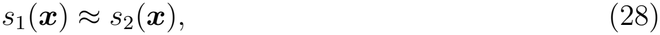

which reveals that, at the module-selection balance, selective benefits provided by adaptive mutations in both modules are identical. In other words, in our simple modular GPFM with a low-entropy optimum, the module-selection balance is achieved on the line of equal fitness benefits in the trait space. Thus, a population adapting on the modular GPFM initially converges to the line of equal fitness benefits and stays in this module-selection balance until it eventually approaches the mutation-selection equilibrium near the fitness peak (Hartl and Taubes, 1998; Sella, 2009; Nourmohammad et al., 2013).

These simulations and analytical results reveal how a tug-of-war between selection and mutational pressures shapes trait evolution in the successive mutations regime. When the fitness optimum is entropically favored, mutational biases are weak, and selection pushes the populations up the fitness gradient. When the fitness optimum is entropically disfavored, mutational biases oppose selection, particularly near the boundaries of the trait space, which in our case is defined by the rays *x*_*i*_ = 0, *i* = 1, 2. On modular GPFM, the mutational pressures are so strong that they invert the curvature of the trait trajectories relative to the gradient (compare trajectories starting at the rhombus between panels B and F in Figure 2) and lead to the emergence of the module-selection balance. The pleiotropic GPFM with low-entropy optimum represents an intermediate case between these two extremes. Here, mutational pressures prevent populations from following the fitness gradient, but they are not strong enough to invert the curvature of trait trajectories and cause convergence to a unique module-selection balance.

### 3.2 Convergence to module-selection balance is enhanced by clonal interference and weakened by sex in the concurrent mutations regime

So far, we observed that the evolutionary dynamics of functional traits change qualitatively in the successive mutations regime depending on the pleiotropic versus modular GPFM architecture and on the entropy of the trait space. We next sought to understand how these dynamics might be affected by the increased supply of adaptive mutations and the mode of reproduction. When the supply of adaptive mutations is sufficiently large, populations transition to the so-called “concurrent mutations” regime where multiple beneficial alleles simultaneously segregate in the population (Desai and Fisher, 2007). In this regime, it becomes important whether or not individuals are able to exchange genetic material (Muller, 1932; Neher and Shraiman, 2009). In this section, we consider trait evolution in the concurrent mutations regime in two extreme cases with respect to reassortment of the two chromosomes, either when they are completely linked (*ρ* = 0) or when they reassort freely (*ρ* = 1). Since the module-selection balance observed in the previous section on the modular GPFM with a low-entropy optimum is an interesting and, to our knowledge, novel phenomenon, we pay particular attention to whether and how convergence to this state changes in the concurrent mutations regime.

#### Concurrent mutations regime, complete linkage

We carried out simulations of the Wright-Fisher model in the concurrent mutations regime with complete linkage between chromosomes (see Section 2.4 in “Methods” for details). We find that, on the pleiotropic GPFM with either a low- or a high-entropy optimum, the dynamics of adaptation in the trait space remain similar to the successive mutations regime (Figure 3). Specifically, on the pleiotropic GPFM with a high-entropy optimum, populations closely follow the paths of steepest ascent (equation (23)), approaching the fitness peak along the *x*-axis, which corresponds to the functional trait 1 that is under weaker selection (Figure 3A). Correspondingly, the module performance ratio decreases over time (Figure 3C). On the pleiotropic GPFM with a low-entropy optimum, populations continue to evolve along relatively straight trajectories (Figure 3E), though a weak convergence trend is evident in the module performance ratio (Figure 3G).

In contrast, the trait dynamics on modular GPFMs are substantially different in the concurrent mutations regime with complete linkage compared to the successive mutations regime. Populations evolving on the modular GPFM with a low-entropy optimum rapidly converge to a module-selection balance defined by equation (28), which describes the line of equal fitness benefits (see Section 2.3.3 in “Methods“). While in this balance, populations approach the fitness peak by improving both functional traits at proportional rates. We derive an analytical approximation for the evolutionary dynamics in the trait space in this regime, which matches our simulated results reasonably well (see Figure 3F, equation (17) in “Methods” and Section 2.2 in Supplementary Information). In particular, it confirms our visual observation that, prior to converging to the line of equal fitness benefits, populations experience evolutionary stalling of the module that is better adapted, consistent with previous studies (Venkataram et al., 2020; Gomez et al., 2020).

The dynamics of populations evolving on the modular GPFM with a high-entropy optimum are qualitatively different in the concurrent mutations regime with complete linkage compared to the successive mutations regime. In contrast to what we have seen in the previous section, here we find that populations no longer follow the fitness gradient and instead appear to also converge to a module-selection balance (Figures 3B,D), albeit one that is different from the line of equal fitness benefits. This convergence is most pronounced in populations with an initially poorly performing functional module 1 (rhombus in Figures 3B). In such populations, the curvature of the average trait trajectories has the opposite sign relative to the trajectories of steepest ascent (compare Figure 3B and Figure 2B).

#### Concurrent mutations regime, unlinked chromosomes

We next examined how reassortment between chromosomes affects the dynamics of trait evolution. The trait trajectories on both pleiotropic GPFMs are unaffected by reassortment (compare Figure 4A,C,E,G with Figure 3A,C,E,G), as expected since the two chromosomes are statistically equivalent in these models. In contrast, reassortment plays an important role on modular GPFMs. We find that the evolution of sexual populations on the modular GPFM with a high-entropy optimum is very similar to the successive mutations regime, i.e., these populations closely follow the fitness gradient (compare Figure 4B with Figure 2B), and correspondingly, the performance ratio steadily decreases (Figure 4D). On the other hand, populations evolving on the modular GPFM with a low-entropy optimum still approach the module-selection balance, but do so more slowly than in the linked case (compare Figures 4H and 3H). A simple analytical approximation based on the Desai-Fisher theory (Desai and Fisher, 2007) and the assumption that the two traits evolve independently (see Section 2.3.3 in “Methods“) matches these mean trajectories reasonably well (Figure 4F).

Taken together, these results confirm that populations traverse the trait space in qualitatively different ways as they adapt on modular versus pleiotropic GPFMs with different distributions of genotypes across the trait space, whether adaptation happens in the successive or concurrent mutations regime. These results also reveal how reassortment between the chromosomes that encode the two traits alters these dynamics on modular GPFMs. We find that strengthening the linkage between chromosomes enhances convergence to a module-selection balance; conversely, increasing the reassortment rate weakens this convergence or even causes the module-selection balance to disappear altogether.

### 3.3 Module-selection balance in the “nested” Fisher’s geometric model

Perhaps the most interesting observation made so far is the emergence of the quasisteady state, which we termed “module-selection balance”. This state emerges on modular GPFMs, particularly when fitness optimum is located in a sparsely populated region of the genotype space. However, since our models are built on several simplifying assumptions, we next sought to better understand the robustness of the module-selection balance with respect to some of them.

To this end, we examine a more complex model that we refer to as the “nested” Fisher’s geometric model (nFGM, Figure 5A). In this model, each “high-level” functional trait *x*_*i*_ is itself decomposed into *n*_*i*_ “low-level” additive functional traits 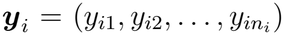 . Specifically, we model *x*_*i*_ as a quadratic function of ***y***_*i*_, 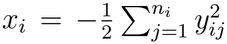 . We assume that the functional modules themselves are tightly integrated, such that the map from genotypes to low-level phenotypes ***y***_*i*_ is highly pleiotropic, and we assume that the optimal phenotype ***y***_*i*_ = **0** is located in the genotype-dense region of the trait space, analogous to the pleiotropic genotype-phenotype map with a high-entropy optimum described above (Figure 1A and Section 2.1 in “Methods“). More specifically, each mutation affecting trait *x*_*i*_ is a vector ***m***_*i*_ in the *n*_*i*_-dimensional space with a fixed magnitude 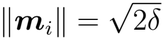 and a random, uniformly distributed direction (see Section 2.1.6 in “Methods” for details). In this model, the effect _Δ*i*_ of a new mutation on the functional trait *x*_*i*_ is approximately normally distributed with mean −*δ* and standard deviation 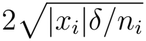 (Tenaillon, 2014).

**Figure 5.**
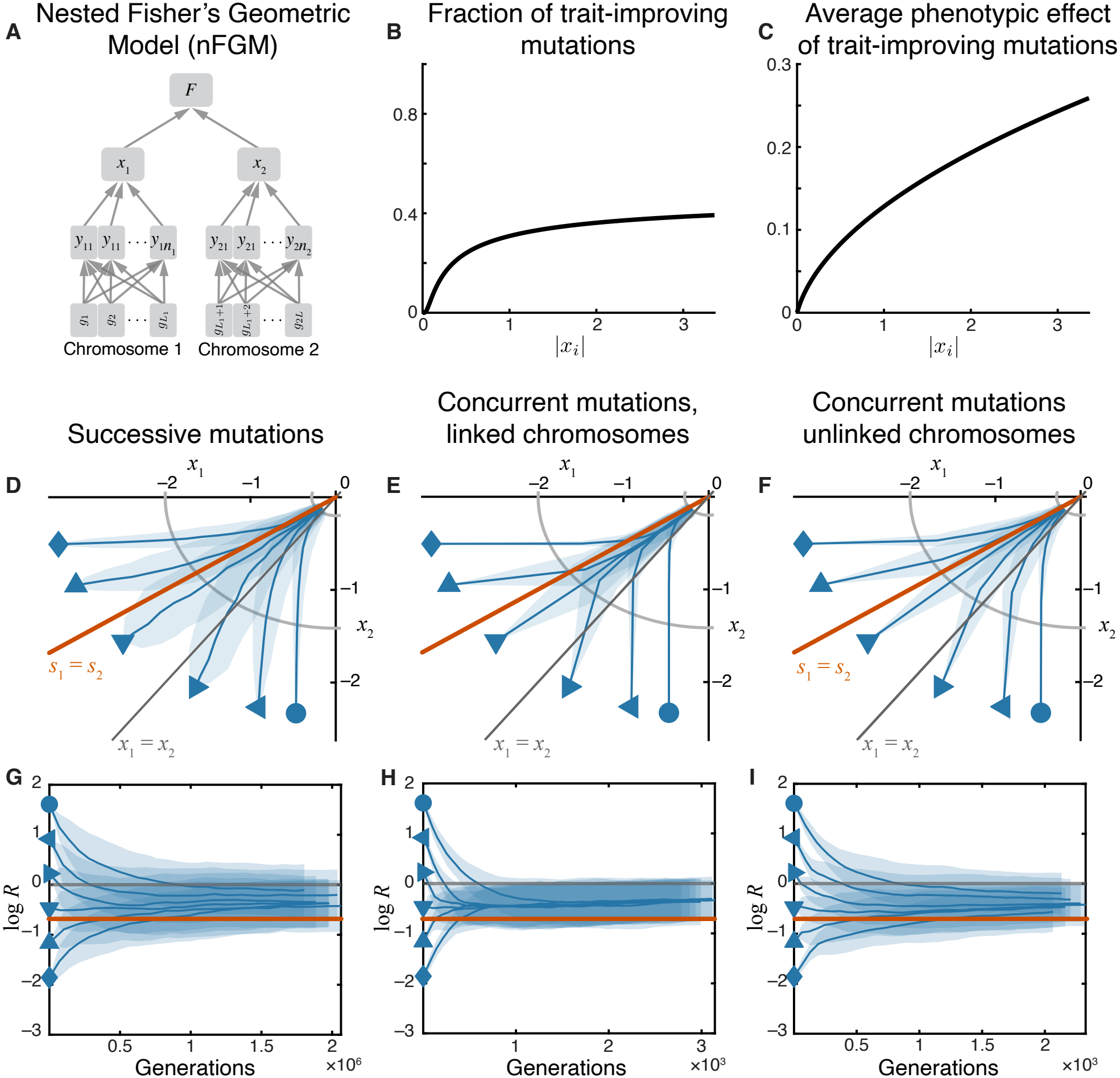
Trait evolution on the nested Fisher’s geometric model (nFGM). A. Schematic of the nFGM. **B**. Fraction of trait-improving mutations as a function of the distance of the trait to its optimal value |*x*_*i*_|. **C**. Average phenotypic effect of a trait-improving mutation as a function of the distance of the trait to its optimal value |*x*_*i*_|. **D–I**. Evolutionary trajectories in the trait space (D–F) and the corresponding module-performance ratios *R* = *x*_2_*/x*_1_ (G–I) for three evolutionary regimes, as indicated, analogous to Figures 2–4. Parameter values: number of lower-level traits per high-level trait is *n*_1_ = *n*_2_ = 10; magnitude of a mutation in the space of lower-level traits is 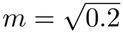 ; genome-wide mutation rate is *U* = 10^*−*7^ in the successive mutations regime and *U* = 2 × 10^*−*4^ in the concurrent mutations regimes; other parameters are the same as in Figures 2–4. See Section 2.1.6 in “Methods” for details.

The fact that the high-level traits are quadratic functions of the low-level traits introduces some important differences between the nFGM model and the simpler models considered in Sections 3.1, 3.2. First, although the density of genotypes in the vicinity of the optimum ***y***_*i*_ = **0** is high, almost all of them have suboptimal values of the high-level trait *x*_*i*_. Therefore, the fitness optimum does not coincide with the peak of genotype density. As a result, the supply of mutations that improve the performance of module *i* declines with its increasing performance (i.e., when *x*_*i*_ is closer to zero). In this sense, the nFGM model is similar to the modular GPFM with a low-entropy optimum considered in Sections 3.1, 3.2. The second consequence of the non-linearity in the mapping ***y***_*i*_ ↦→ *x*_*i*_ is that the high-level traits *x*_*i*_ are non-additive, which implies that both the supply of mutations that improve module *i* and the distribution of their effect sizes are non-linear functions of *x*_*i*_ (Figure 5B,C). Therefore, we can think of nFGM as probing the robustness of the module-selection balance with respect to the assumption of trait additivity and the assumption that all mutations have the same and constant phenotypic effect *δ*.

We simulated evolution on the nFGM model with ten-dimensional low-level traits (*n*_1_ = *n*_2_ = 10) in the same three evolutionary regimes as before. We found that populations approach a module-selection balance (Figure 5) and that the convergence to this quasi-steady state is even more apparent than in our simple modular GPFM. Interestingly, when both modules have the same underlying dimensionality *n*_*i*_, the module performance ratio equilibrates close to the line of equal fitness benefits given by equation (26) which is predicted by our simple modular GPFM model with a low-entropy optimum (Figure 5). When the underlying module dimensionalities are different, module-selection balance is still achieved, albeit at a different equilibrium performance ratio (Figure S5). These results suggest that the existence of a module-selection balance is robust with respect to the precise functional form of how the supply and the effects of module-improving mutations depend on the distance to the phenotypic optimum.

### 3.4 Patterns of genome evolution in Lenski’s LTEE are consistent with an approach to a module-selection balance

The simple GPFMs that we have analyzed in this work are but caricatures of real biological GPFMs. However, if they capture some essential properties of biological GPFMs, our results should give us an insight into the evolutionary dynamics of empirical populations. As mentioned above, there is substantial evidence that biological GPFMs are variationally modular rather than highly pleiotropic (Wagner et al., 2007; Wagner and Zhang, 2011). Although much less is known about how genotypes are distributed in the relevant trait spaces, biophysical models and experimental studies show that functional proteins are encoded by highly non-random DNA sequences (Zou and Saven, 2000; Keefe and Szostak, 2001; Taverna and Goldstein, 2002; Koehl and Levitt, 2002; Sinai et al., 2026), which suggests that the entropy of trait values conferring high fitness is probably quite low. Thus, among our simple GPFM models, the modular GPFM with a low-entropy optimum may bear the closest resemblance to biological GPFMs.

One salient feature of evolution on the modular GPFM with a low-entropy optimum is that trait trajectories are often bi-phasic, particularly in the concurrent mutations regime with complete linkage (Figure 3F). The initial phase of adaptation is characterized by evolutionary stalling whereby the worse-performing module improves while the better-performing module does not (Venkataram et al., 2020). In the second phase, populations are in the module-selection balance where both functional modules improve at proportional rates. Notably, we observed similar bi-phasic trajectories even when the optimum was entropically favored, as long as populations were adapting in the concurrent mutations regime with complete linkage (Figure 3B). In contrast, evolution on pleiotropic GPFMs is characterized by simultaneous improvement of both modules along the entire course of evolution and lacks any obvious bi-phasic structure (Figure 3A,E). Given this contrast and strong indications that biological GPFMs are variationally modular, we wondered whether we might be able to detect signatures of bi-phasic trait evolution in asexual experimental populations evolving for a long time in a constant environment in the concurrent mutations regime.

Lenski’s long-term evolution experiment (LTEE) is probably the best system to address this question (Lenski et al., 1991; Lenski, 2017). In this experiment, 12 replicate populations of the bacterium *Escherichia coli* have been evolving asexually in a constant laboratory environment in the concurrent mutations regime over many thousands of generations (Good et al., 2017). We would ideally like to examine long-term quantitative temporally resolved changes in functional traits relevant for fitness in these populations, but such data are not readily available. Thus, instead, we turn to published whole-genome time-course sequencing data from these populations (Good et al., 2017).

Any examination of LTEE genomic data through the lense of our theory faces two challenges. First, our models do not explicitly describe patterns of genome evolution. And second, we have only considered GPFMs with two functional traits, whereas the number of traits relevant for fitness in *E*.*coli* is likely larger, although some studies suggest that it is quite small, perhaps less than ten (Kinsler et al., 2020; Iwasawa et al., 2022; Donyavi et al., 2025; Ghosh et al., 2026). How populations would traverse trait spaces on GPFMs with more than two dimensions is currently unclear. Thus, before examining the LTEE data, we develop some basic expectations for what patterns we might expect at the genome level in organisms with multiple traits.

#### Expectations for genome evolution on modular GPFMs

We first ask what patterns of genome evolution we might expect on variationally modular GPFMs with two functional traits (Figure 1B). Since different functional traits are encoded in different parts of the genome (i.e., by distinct genes), the evolutionary stalling of one trait in the initial phases of adaptation (Figure 3B,F) implies that mutations that drive this early adaptation are concentrated in a subset of genes, those that encode the improving trait. In contrast, once the population achieves a module-selection balance where all modules are improving (Figure 3B,F), mutations that drive adaptation are distributed across genes that encode both traits.

Extrapolating this reasoning to organisms with more than two functional traits, we would expect that, if one or a few traits are much more suboptimal than others in the starting population, natural selection would initially improve these traits while the remaining better-performing traits would be stalled. At later stages of evolution, we would expect populations to approach a module-selection balance where all traits improve simultaneously. Thus, we expect the distribution of driver mutations across the genome to follow qualitatively similar patterns on modular GPFMs with higher trait dimensionality: driver mutations should be concentrated in relatively few genes at the early stages of adaptation, and distributed more broadly at later stages of adaptation.

To test this intuition, we simulated the evolution of organisms with 10 functional traits in the concurrent mutations regime with complete linkage (see Section 4 in Supplementary Information for details). As expected, we found that, as long as some modules initially performed substantially worse than others, early evolutionary improvements were concentrated on a subset of modules, consistent with evolutionary stalling (Figure S6D,H and Figure S7A). At later stages of evolution, improvements were distributed more uniformly across all modules, consistent with the approach to a module-selection balance (Figure S6D,H). These patterns held for modular GPFMs with either high- or low-entropy optima.

#### Expectations for genome evolution on pleiotropic GPFMs

As discussed in previous sections, populations adapting on pleiotropic GPFMs traverse the trait space differently than populations evolving on modular GPFMs, particularly in the concurrent mutations regime with full linkage (Figure 3). While the specific shape of trait evolutionary trajectories depends on the distribution of genotypes in the trait space, the salient feature of adaptation on two-trait pleiotropic GPFMs is that both traits improve simultaneously throughout the course of evolution (Figure 3A,E). Our simulations show that this remains true in higher-dimensional trait spaces as well (Figure S6A,C,E,G).

Deriving clear-cut expectations for the genomic distribution of driver mutations underpinning such trajectories is more difficult for pleiotropic GPFMs because these distributions depend on the association between the mutation’s degree of pleiotropy (represented by the angle *θ*_*l*_ in equation (2)) and its genomic location. In our simple models, where no such association exists, the genomic distribution of driver mutations would remain close to uniform throughout the course of adaptation. If the degree of pleiotropy is associated with the genomic location (i.e., mutations with certain values of angle *θ*_*l*_ are more likely to be found in certain genes), the distribution of driver mutations along the genome could change over time, but there is no generic reason to expect that this distribution should broaden or become narrower as adaptation proceeds.

This reasoning suggests that the main distinctive feature of genome evolution on modular GPFMs is that the breadth of the distribution of driver mutations along the genome should increase over time. Thus, we next looked for this pattern in the genomic data from Lenski’s LTEE.

#### Patterns of genome evolution in Lenski’s LTEE

We reanalyzed genomic data from six non-mutator LTEE populations obtained by Good et al. (2017). Across these populations, Good et al. (2017) identified 1,670 mutations that passed their filters. We focused our analysis on 1,464 of these mutations that fall within annotated genes (see Section 2.5 in “Methods“). Good et al. (2017) estimated the detection time for each mutation and found that these times are non-randomly distributed, which suggests that the targets of natural selection shifted over time. In particular, they reported a significant enrichment of parallel mutations before generation 17,500.

To examine how the breadth of the genomic distribution of adaptive mutations changes over time, we computed the effective number of genes targeted by selection in 10^4^-generation sliding windows (see Section 2.5 in “Methods“). We find that this number rises sharply from ∼55 at the beginning of the experiment to ∼86 by roughly 17,500 generations, after which it remains approximately constant (Figure 6A). This pattern is also observed among mutations that occurred in multi-hit genes, which are strongly enriched for driver mutations (Figure S8). A direct visualization of the distribution of mutations along the genome reveals a clear clustering of early mutations at certain genomic loci and a more uniform distribution of late mutations (Figure 6B). This bi-phasic pattern is consistent with LTEE populations evolving on a variationally modular GPFM and reaching a module-selection balance after about ∼ 17,500 generations.

**Figure 6.**
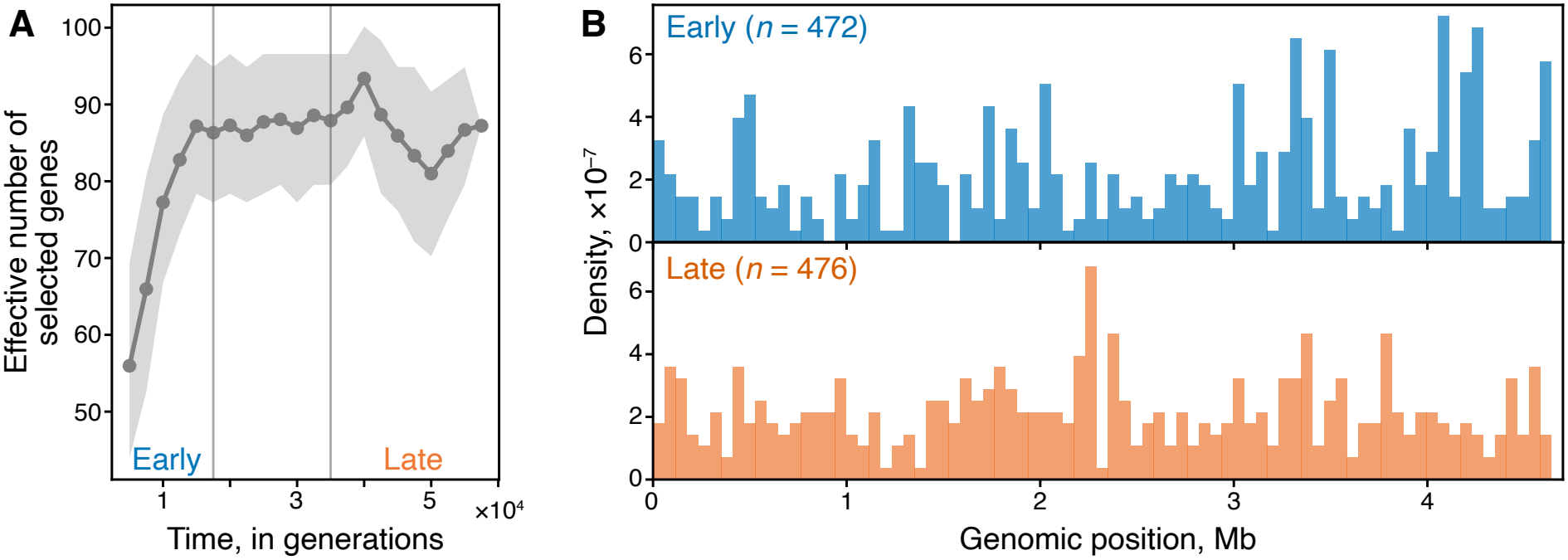
Distribution of mutations across the *E. coli* genome in Lenski’s LTEE. A. Effective number of genes targeted by natural selection (computed as the reciprocal of the Simpson’s index) over 10^4^-generation sliding windows. The thick line shows the reciprocal of Simpson’s index averaged over 2 × 10^3^ subsamples of 104 mutations drawn without replacement, and the shaded region spans the 2.5th to 97.5th percentiles across subsamples (see Section 2.5 for details). **B**. Genomic density of mutations that arose in the early (first 17,500 generations; *n* = 472 mutations) and in the late epoch (final 25,500 generations; *n* = 476 mutations). Data are from six non-mutator LTEE populations sequenced by Good et al. (2017).

## 4 Discussion

Here, we used several modifications and extensions of the two-dimensional Fisher’s geometric model to investigate how adapting populations traverse the trait space in a constant environment. We examined how these trajectories are affected by two factors, the degree of variational modularity of the genotype-phenotype map and the distribution of geno-types in the trait space. We considered four simple GPFM models at the extremes of a continuum with respect to each factor: a variationally modular GPFM and a GPFM with universal pleiotropy where the optimal trait values are located in a high- and a low-entropy regions of the trait space (Figure 1). We investigated the mean trait evolutionary trajectories on these GPFMs in three regimes of evolutionary dynamics: the successive mutations regime (also known as the strong-selection, weak mutation regime), the concurrent mutations regime with asexual reproduction, where the genome is completely linked, and the concurrent mutations regime with sexual reproduction, where the chromosomes encoding different functional traits reassort at each reproduction event.

We found that, on the highly pleiotropic GPFMs, the average evolutionary trajectories in the trait space remain similar across the regimes of evolutionary dynamics. On the GPFM with universal pleiotropy and a high-entropy optimum (Figure 1A,C), an average population adapts by following the fitness gradient (Panel A in Figures 2–4). As a result, the functional module under stronger selection approaches its optimum exponentially faster than the functional module under weaker selection, which yields a continually declining ratio of module performances (Panel C in Figures 2–4). On the pleiotropic GPFM with a low-entropy optimum (Figure 1A,E), populations move towards the fitness optimum along relatively straight lines in the trait space (Panel E in Figures 2–4). Correspondingly, the ratio of module performances remains nearly constant, although we observe some weak signs of convergence to an intermediate ratio in populations evolving in the concurrent mutations regime (Panel G in Figures 2–4).

On the other hand, on modular GPFMs, trait trajectories vary significantly across the regimes of evolutionary dynamics. On the modular GPFM with a low-entropy optimum (Figure 1B,F), populations evolving in any of the three regimes converge to a quasi-steady state, which we termed the “module-selection balance”. In this state, both modules improve at proportional rates and their performance ratio remains constant over time as they approach the fitness peak (Panels F and H in Figures 2–4). In the simple GPFM models considered here, the module-selection balance is achieved on the line of equal fitness benefits where the selection coefficients of mutations improving each trait are equal. Although populations converge to a module-selection balance in all regimes of evolutionary dynamics, the shape of trait trajectories differs markedly across regimes. In particular, we found that asexual populations evolving in the concurrent mutations regime converge to the line of equal fitness benefits along fairly straight paths, i.e., if one trait is initially much farther from its optimum than the other, adaptation occurs almost exclusively by improving the lagging trait whereas the more optimal trait is stalled (Figure 3F). In contrast, sexual populations or those evolving in the successive mutations regime converge to a module-selection balance more gradually without exhibiting evolutionary stalling (Panel F in Figures 2 and 4).

Finally, the trait evolutionary trajectories that emerge on the modular GPFM with a high-entropy optimum (Figure 1B,D) display a mixture of features observed on other GPFMs. Populations that evolve on this GPFM either in the successive mutations regime or in the concurrent mutations regime with sexual reproduction traverse the trait space along the paths of steepest ascent (Panel B in Figures 2 and 4), similar to the pleiotropic GPFM with a high-entropy optimum, whereas asexual populations evolving in the concurrent mutations regime converge to a module-selection balance (Figure 3B), similar to the modular GPFM with a low-entropy optimum.

Our results demonstrate how biases in the phenotypic effects of mutations interact with selection in shaping the adaptive trajectories of populations in their trait spaces. Broadly, selection pushes populations along the gradient of the fitness surface, but mutational biases constrain how closely populations can follow this path of steepest ascent. The role of mutational biases in evolutionary dynamics has received significant attention in recent literature (Yampolsky and Stoltzfus, 2001; Stoltzfus and McCandlish, 2017; Storz et al., 2019; Gomez et al., 2020; Cano and Payne, 2020; Cano et al., 2022; Sane et al., 2023; Tuffaha et al., 2023; Gitschlag et al., 2023; Parveen et al., 2025; Fuqua and Vakirlis, 2026; Barber and Couce, 2026), but most of these studies have focused on biases that arise from the mutational process, such as how often transversions happen compared to transitions. In contrast, here we investigated mutational biases that arise generically at the phenotypic level due to the genetic encoding of traits and the distribution of genotypes in the trait space. Biases of this sort are expected to arise in any kind of genotype-phenotype map, even if all types of mutations at the genetic level occur at the same rate. Thus, our work is closely related to the discussion of the role of developmental constraints in evolution (e.g. Maynard Smith et al., 1985; Klingenberg, 2005; Brakefield, 2006; Losos, 2011).

The intuition behind how selection and mutation jointly shape the evolutionary trajectories in the trait space can be most easily gained in the successive mutations regime (Figure 2). On the pleiotropic GPFM with a high-entropy optimum, mutational biases are nearly absent in the vicinity of the optimum, i.e., all mutational directions are available and nearly equiprobable (Figure 1C). Therefore, selection is the only remaining force, and populations follow the trajectories of steepest ascent. Variational modularity restricts the availability of mutations and forces selection to “choose” whether to fix a mutation that improves trait 1 or trait 2. Consider for example a situation when trait 2 is much closer to its optimal value than trait 1 (rhombus in Figure 2F). Here, the fitness gradient still has a significant component along the *x*_2_ axis, but since improving the lagging trait 1 yields a higher fitness gain, selection more often “chooses” such mutations over those that improve only trait 2. Interestingly, the lack of pleiotropic mutations is on its own not sufficient to distort the average trajectories away from the path of steepest ascent. If both types of mutations are equally available, as is the case on the GPFM with a high-entropy optimum (Figure 1D), selection fixes either of them with the probability proportional to its fitness effects, such that the average population continues to move along the fitness gradient (Figure 2B). In contrast, if the optimum is located in a low-entropy region of the trait space (Figure 1F), mutations improving the lagging trait are not only favored by selection, they are also more readily available. As a result, such mutations fix more often, and the curvature of trait trajectories inverts relative to the paths of steepest ascent (compare trajectories originating at the rhombus in panels B and F in Figure 2).

The pleiotropic GPFM with a low-entropy optimum is an interesting intermediate case that illustrates the role of trait-space entropy in the absence of variational modularity. Despite the highly pleiotropic genetic architecture, the entropy alone can introduce strong biases in the availability of mutations. For concreteness, consider again a situation when trait 2 is much closer to its optimal value than trait 1 (rhombus in Figure 2E). Such a trait combination can only be realized by very atypical genotypes, which have access to atypical distributions of mutational effects. In our model, genotypes populating this part of the trait space carry allele 1 only at sites with *θ*_*l*_ ≈ *π*; as a result, they have access to beneficial mutations that are biased towards improving the lagging trait 1. In our model, the magnitude of this bias is sufficiently strong to distort evolutionary trajectories from the path of steepest ascent but not strong enough to cause convergence to a module-selection balance. However, our model is just one particular example of how trait space may be populated by genotypes, and whether such nearly linear trait trajectories are a general feature of highly pleiotropic GPFMs with low-entropy optima is unclear. Further work is need to better understand the mutational biases and the evolutionary dynamics arising on such GPFMs.

Building upon the above discussion of the concurrent mutations regime, we can begin developing an intuitive understanding of why trait trajectories on modular GPFMs are affected by clonal interference and sexual reproduction. In asexual populations evolving in the concurrent mutations regime, clonal interference prevents the fixation of all but the most beneficial mutations (Strelkowa and Lässig, 2012; Good et al., 2012). Since mutations that improve the lagging trait provide larger fitness benefits than mutations improving the already more optimal trait, the former are fixed preferentially and the population tends to move towards the interior of the trait space, even on GPFMs with a high-entropy optimum (Figure 3B). Entropy-induced mutational biases further amplify this trend (Figure 3F). In contrast, reassortment between chromosomes that encode different traits creates high-fitness genotypes that carry combinations of beneficial alleles in both traits. Thus, sexual reproduction alleviates the mutational biases imposed by modularity. On the modular GPFM with a high-entropy optimum, this enables populations to follow the fitness gradient (Figure 4B) and on the modular GPFM with a low-entropy optimum, it weakens convergence to a module-selection balance (Figure 4F).

The module-selection balance observed on modular GPFMs, particularly ones with a low-entropy optimum, is an interesting phenomenon that, to our knowledge, has not been described in the prior literature. This begs the question: do biological GPFMs look anything like our simple modular GPFM with a low-entropy optimum? Although we cannot be certain at this point, we believe that the existing evidence points in the right direction. Many studies have shown that the effects of genetic perturbations are confined to subsets of traits, i.e., pleiotropy is far from “universal” (Wagner and Zhang, 2011). Furthermore, finding functional proteins among random amino acid sequences appears to be difficult (Keefe and Szostak, 2001), which suggests that fitness optima are located in low-rather than high-entropy regions of the trait space, at least for molecular phenotypes.

We discovered the module-selection balance in a relatively simple GPFM model with two traits, but found that it is preserved and even strengthened in more complex models with non-additive traits or higher-dimensional trait spaces. The fact that convergence to a module-selection balance is preserved in these more complex models suggests that this quasi-steady state may be a generic feature of long-term evolution on variationally modular GPFMs, particularly those with low-entropy optima, and especially in large asexual populations, which tend to evolve in the concurrent mutations regime.

As discussed in the Introduction, our work was in part motivated by the empirical observation of evolutionary stalling (Venkataram et al., 2020) and the theoretical explanation behind it (Gomez et al., 2020), i.e., the fact that, in the presence of clonal interference, natural selection focuses on improving the lagging module while the leading one is stalled despite the availability of beneficial mutations in it. Our work places these previous results into a broader dynamical context. In particular, as hypothesized by Venkataram et al. (2020), we have shown that evolutionary stalling is a transient phase of adaptation that is supplanted by a module-selection balance where both modules improve at proportional rates.

In so far as our simple GPFM models capture some essential features of GPFMs of biological organisms—most importantly, their variational modularity and, to a lesser degree, the trait-space entropy—we would expect rapidly adapting asexual populations to exhibit similar bi-phasic dynamics as we observed in these models (Figures 3B,F, 5E, S6D,H, S7A). Specifically, if there is a large disparity in the initial performance of functional modules, natural selection is expected to be initially focused on improving modules whose performance is most sub-optimal, leading to a relatively narrow distribution of adaptive mutations along the genome. Once the population approaches a module-selection balance, our models suggest that many modules should improve simultaneously, leading to a relatively broad distribution of adaptive mutations along the genome. In other words, we expect the genomic distribution of adaptive mutations to broaden over time.

We tested this prediction in Lenski’s LTEE using metagenomic data obtained by Good et al. (2017) and found a strikingly bi-phasic change in the genomic distribution of putatively adaptive mutations, consistent with our model (Figure 6). The abrupt shift in the width of the distribution of adaptive mutations occurs between 15 and 20 thousand generations. Notably, fitness gains in LTEE populations dramatically slow down at approximately the same time, an observation that gave rise to the “two-epoch” hypothesis proposed earlier by Good and Desai (2015). It is possible that the first epoch, characterized by rapid fitness gains, corresponds to evolutionary stalling of some modules, whereas the second epoch, characterized by much slower fitness gains, corresponds to the module-selection balance.

The fact that the patterns in the genomic data from Lenski’s LTEE are consistent with an approach to a module-selection balance may appear trivial at first. After all, GPFMs of most if not all organisms are believed to be at least to some extent variationally modular (Wagner and Zhang, 2011). Instead, the significance of this finding is threefold. First, it suggests that a module-selection balance—which was derived in a simple model with just two orthogonal traits and perfect variational modularity—can in fact be achieved even on a potentially very complex GPFM with many “fuzzy” modules. Second, convergence to a quasi-steady state implies that any phenotypic differences that may have been present between initial populations is erased. If biological populations indeed converge to a module-selection balance so rapidly (within about 10^4^ generations for moderately large populations with *N*_*e*_ ≈ 3.3 × 10^7^ (Lenski et al., 1991)), it suggests that a relatively short bout of adaptation in a constant environment is sufficient to erase phenotypic differences between populations long before these populations approach the fitness peak. Perhaps most importantly, module-selection balance implies that, even if the underlying trait space is high-dimensional, populations might approach fitness peaks along low-dimensional manifolds. Such manifolds are characterized by certain phenotypic constraints and constraints on the fitness effects of beneficial mutations. For example, in our simple models, module-selection balance is achieved on the line of equal fitness benefits where all beneficial mutations have the same selection coefficients (in a given genetic background). This suggests that well-adapted organisms may admit particularly simple effective descriptions, consistent with recent observations of “emergent simplicity” (e.g. Bergelson et al., 2021; Kinsler et al., 2020; Scott and Hwa, 2023; Ardell et al., 2024).

## 5 Acknowledgments

We thank the Kryazhimskiy lab, the Meyer labs and Matt Pennell for feedback on the project, Jhelam Deshpande for comments on an early version of the manuscript. This work was supported by the NIH grants 1R01GM137112 and R35GM153242 to SK. SMA was supported by the NIH T32 program (5T32GM133351-02) and the Curci Foundation.

