## Supplementary Information for "Module-Selection Balance in the Evolution of Modular Organisms"

**5 Supplementary Figures** **14**

### 1 A note on the definition of "variational modularity"

In an authoritative review, Wagner et al. (2007) define a variational module as "*a set of covarying traits that vary relatively independently of other such sets of traits*" and add that "*variational modules are recognized by higher than average correlations among traits*". This is a general definition, and, as such, it leaves some room for interpretation. For example, how the variation is generated in the first place is unspecified, which means that variational modules identified with respect to different ensembles could be different, e.g., traits with zero correlation among single-mutation neighbors of a genotype may be highly correlated among adult individuals in a wild population. In this work, we are interested in the entire genotype-phenotype map, i.e., our ensemble is the space of all genotypes.

Another important ambiguity arises from the meaning of trait independence. One natural interpretation of independence is *statistical*, as the absence of correlations among certain traits in the sample. Another natural interpretation is *genetic*, as different traits being encoded in different parts of the genome. It appears that different communities of evolutionary biologists have adopted somewhat different conventions on how to resolve this ambiguity. Specifically, in the quantitative genetics literature, the phenotypic effects of mutations are typically modeled as random draws from multivariate normal distributions. In this context, traits independence is naturally interpreted in the statistical sense as zero value of the respective covariance term (e.g., Pavlicev et al., 2010; Melo and Marroig, 2015; Chebib and Guillaume, 2022; do O and Whitlock, 2023). In contrast, papers that fall more into the domain of population genetics, typically interpret independence genetically as each mutation affecting only one or a subset of traits (e.g., Welch and Waxman, 2003; Gutenkunst and Sethna, 2007). It is this latter convention that we have adopted in this work (Figure 1B in the main text).

One can immediately notice that the genetic interpretation of independence is more stringent, in the sense that traits that are encoded in different parts of the genome must also be statistically independent, but the converse is not true. If we consider the entire genotype space, a genotype-phenotype map that is modular according to the genetic interpretation is also modular according to the statistical interpretation, but a genotype-phenotype map that is modular in the statistical sense may not be modular in the genetic sense (e.g. if each allele flip has effects on both traits but the magnitudes of these effects themselves are uncorrelated, as in our pleiotropic GPFM with a high-entropy optimum, see Section 2.3.1 in the main text).

### 2 Evolutionary dynamics on the modular GPFM with a low-entropy optimum in the concurrent mutations regime with complete linkage

#### 2.1 Numerical confirmation of the results by Gomez et al. (2020)

Gomez et al. (2020) studied adaptation of an asexual population of an organism with two traits in the concurrent mutations regime. In their model, each trait $k$ is characterized by its own selection coefficient $s_k$ and beneficial mutation rate $U_{bk}$. Their central finding is that adaptation of the two-trait organism can be understood by studying adaptation of two respective single-trait organisms. We refer to the rate of adaptation of an organism with a single trait $k$ by $v'_k$ and say that $v'_k$ is the rate of adaptation of trait $k$ "in isolation". And we refer to the rate of adaptation of a two-trait organism due to mutations improving trait $k$ by $v_k$ and say that $v_k$ is the rate of adaptation of trait $k$.

Gomez et al. (2020) show that the rate of adaptation $v_k$ of trait $k$ in a two-trait organism depends primarily on whether that trait has a faster or slower rate of adaptation in isolation $v'_k$, and not on the specific combination of $s_k$ and $U_{bk}$ that produces $v'_k$. In particular, if one trait $j$ would adapt much faster than the other trait $i$ in isolation ($v'_j \gg v'_i$), then in a two-trait organism, this trait $j$ adapts at roughly the same rate as in isolation ($v_j \approx v'_j$), while the slower trait $i$ stalls ($v_i \lesssim v'_i \ll v'_j$). Conversely, when both traits would adapt at similar rates in isolation ($v'_i \approx v'_j$), they continue to adapt at similar rates in a two-trait organism ($v_i \approx v_j$). Here, we validate and extend these findings with our own simulations with Poisson offspring sampling across a range of parameter combinations.

**Choice of parameters.** We varied parameters $s_1$, $U_{b1}$, $s_2$, $U_{b2}$ as follows. First, we chose the theoretically predicted rates of adaptation in isolation, $v'_1$ and $v'_2$, given by equation (16) in the main text, from 21 unique pairs spaced over a $6 \times 6$ logarithmic grid with $v'_2 \geq v'_1$ (Figure S1A), where the six values are log-spaced from $5 \times 10^{-6}$ to $10^{-2}$ per generation, so that successive values differ by a factor of $2000^{1/5} \approx 4.57$. Then, for each target value of $v'_i$, we identified four distinct parameter combinations $(s_i, U_{bi})$ that yield $v'_i$ by numerically solving equation $v'_i = f_{\rm DF}(s_i, U_{bi}, N)$ with a fixed ratio $s_i/U_{bi}$ which we varied over the range $\{10, 22, 46, 100\}$, so that $s_i/U_{bi} \gg 1$ to ensure that the Desai-Fisher approximation remains valid (Desai and Fisher, 2007), where $f_{\rm DF}$ is given by equation (16) in the main text. The resulting ranges of $(s_i, U_{bi})$ are shown in Figure S1E–F. This procedure generated $4 \times 4 \times 21 = 336$ total parameter combinations $(s_1, U_{b1}, s_2, U_{b2})$.

**Steady-state simulations.** For each parameter combination, we simulated a haploid asexual population of expected size $N$ with two traits, each subject to beneficial mutations that combine additively. Specifically, at each generation, the fitness of each lineage

$h$ is computed from its accumulated mutation counts $(\nu_{h1}, \nu_{h2})$ as $F_h = \nu_{h1}s_1 + \nu_{h2}s_2$. If lineage $h$ contains $c_h$ individuals, its offspring count is drawn independently from a Poisson distribution with mean $Nc_h e^{F_h} / \sum_j c_j e^{F_j}$. Thus the total offspring population fluctuates, with expected size $N$. Mutations are assigned after the new generation is formed. Each offspring receives a mutation with probability $U_{b1} + U_{b2}$, and the mutation affects trait 1 with probability $U_{b1}/(U_{b1} + U_{b2})$ and trait 2 otherwise. We ran our simulations for a total of $1.5 \times 10^4$ generations with a burn-in of $10^4$ generations to allow the fitness distribution within the population to reach a steady state. The rate of fitness gain attributable to improvements in trait $i$ was estimated from the change in its population mean fitness component $i$ between generations $t_1 = 10^4$ and $t_2 = 1.5 \times 10^4$ as $v_i = \left(\langle \nu_i s_i \rangle_{t_2} - \langle \nu_i s_i \rangle_{t_1}\right) / (t_2 - t_1)$, where $\langle \cdot \rangle_t$ refers to the average over all lineages within the population at generation $t$. If, on average, fewer than two mutations accumulated in a trait during this interval, this trait was classified as stalled; for visualization purposes, stalled traits were assigned a rate of $3 \times 10^{-7}$ per generation (Figure S1B).

**Results.** Our simulations confirm three central findings of Gomez et al. (2020) that underpin the heuristic approximation developed below.

1. Figures S1A,B show that the rates of adaptation $\boldsymbol{v} = (v_1, v_2)$ in a two-trait organism depend primarily on the rates of adaptation in isolation $\boldsymbol{v'} = (v'_1, v'_2)$, and are largely insensitive to the specific values of $s_i$ and $U_{bi}$ that produce a given $\boldsymbol{v'}$. Indeed, we see that the data points in Figure S1B that correspond to the same theoretical $(v'_1, v'_2)$ pair but different underlying $(s_i, U_{bi})$ combinations cluster tightly together in the $(v_1, v_2)$ plane.

2. Figure S1C shows that, when the two traits have very different rates in isolation $(v'_j/v'_i \gg 1)$, the faster trait improves in a two-trait organism at a rate that is close to its rate in isolation $(v_j \approx v'_j)$, while the slower trait contributes very little to adaptation $(v_i \ll v_j)$, consistent with evolutionary stalling driven by clonal interference (Gomez et al., 2020; Venkataram et al., 2020).

3. Conversely, when the two traits adapt at relatively similar rates in isolation, their rates of improvement in a two-trait organism are also similar (Figure S1D). The heuristic approximation for $v_i$ that we develop below operates in this regime.

#### 2.2 Heuristic approximation for evolutionary dynamics

Based on the results of Gomez et al. (2020) discussed above, we approximate the evolutionary dynamics of a population on the modular GPFM in the concurrent mutations regime with complete linkage using the following piecewise approach. We divide the trait space into three regions. Two regions describe the "stalling regime" where one trait supports a much faster rate of adaptation in isolation than the other, i.e., $v'_j/v'_i \gg 1$. In this regime, we approximate the rates of trait improvements as $r_i = 0$ for the slower trait and $r_j = (\delta/s_j)\, v'_j = (\delta/s_j)\, f_{\mathrm{DF}}(s_j, U_{bj}, N)$ for the faster trait. The third region describes the "simultaneous improvement regime" where the two traits support rates of adaptation in isolation that differ by less than a factor of $D$, i.e., $1/D \leq v'_j/v'_i \leq D$. In this regime, we apply the following heuristic approach for approximating $\boldsymbol{v}$. In general, $v_i$ is an unknown function of five parameters, $v_i = \phi\,(s_i, U_{bi}, s_j, U_{bj}, N)$. However, in the special case when the selection coefficients of mutations in the two modules are the same, $s_1 = s_2 = s$, Gomez et al. (2020) have shown that the rate of fitness gains $v_i$ attributable to mutations in trait $i$ is given by

$$\phi(s, U_{bi}, s, U_{bj}, N) = \frac{U_{bi}}{U_b}\, f_{\mathrm{DF}}\,(s, U_b, N)\,, \tag{S1}$$

where $U_{bi}$ is the rate of beneficial mutations in module $i$ as before and $U_b = U_{b1} + U_{b2}$. Thus, we approximate an organism with access to two types of mutations with different selection coefficients $s_1$ and $s_2$ that arise with rates $U_{b1}$ and $U_{b2}$ by an organism with access to two types of mutations with the same selection coefficient $\tilde{s}$ that arise with effective beneficial mutation rates $\tilde{U}_{b1}$ and $\tilde{U}_{b2}$. We then use the functional form (S1) as an ansatz for $\phi$, i.e., we will look for effective parameters

$$\begin{aligned} \tilde{s} &= \xi(s_1, U_{b1}, s_2, U_{b2}, N), \\ \tilde{U}_{bi} &= \eta(s_i, U_{bi}, s_j, U_{bj}, N), \end{aligned}$$

such that

$$v_i = \phi(s_i, U_{bi}, s_j, U_{bj}, N) \approx \tilde{v}_i \equiv \frac{\tilde{U}_{bi}}{\tilde{U}_b}\, f_{\mathrm{DF}}(\tilde{s}, \tilde{U}_b, N), \tag{S2}$$

where $\tilde{U}_b = \tilde{U}_{b1} + \tilde{U}_{b2}$.

Clearly, function $\xi$ must be symmetric with respect to the relabelling of the two traits, i.e.,

$$\xi(s_1, U_{b1}, s_2, U_{b2}, N) = \xi(s_2, U_{b2}, s_1, U_{b1}, N). \tag{S3}$$

In addition, functions $\xi$, $\eta$ must satisfy the basic constraints

$$\xi(s, U_{b1}, s, U_{b2}, N) \;=\; s, \tag{S4}$$

$$\eta(s, U_{bi}, s, U_{bj}, N) \;=\; U_{bi} \tag{S5}$$

that guarantee that the ansatz recovers the equal-selection-coefficient expression in equation (S1) whenever $s_1 = s_2 = s$.

We first assume that we have already found a suitable value $\tilde{s}$ and discuss the choice of the function $\eta$. We return to the choice of the function $\xi$ below. If $\tilde{s}$ is known, to set the effective beneficial mutation rates $\tilde{U}_{bi}$, $i = 1, 2$, we use the observation by Gomez et al. (2020) that $\boldsymbol{v}$ depends primarily on $\boldsymbol{v}'$. Thus, we need to define $\tilde{U}_{bi}$ so that it preserves the rate of adaptation $v_i'$ of trait $i$ in isolation upon substituting $s_i$ by $\tilde{s}$. This reasoning implies that we can define $\tilde{U}_{bi}$ as a solution of equation

$$f_{\mathrm{DF}}(\tilde{s}, \tilde{U}_{bi}, N) = f_{\mathrm{DF}}\left(s_i, U_{bi}, N\right). \tag{S6}$$

Note that, if $s_1 = s_2 = s$, then, according to the constraint (S4), $\tilde{s} = s$. Therefore, equation (S6) becomes

$$f_{\mathrm{DF}}(s, \tilde{U}_{bi}, N) = f_{\mathrm{DF}}\left(s, U_{bi}, N\right),$$

which implies that the condition (S5) is satisfied.

We can rewrite equation (S6) as

$$v_i'\, z_i^2 + \tilde{s}^2\, z_i - 2\tilde{s}^2 \log(N\tilde{s}) = 0,$$

where $z_i = \log(\tilde{s}/\tilde{U}_{bi})$, $v_i' = f_{\mathrm{DF}}\left(s_i, U_{bi}, N\right)$, and find that

$$z_i = \frac{-\tilde{s}^2 \pm \sqrt{\tilde{s}^4 + 8\tilde{s}^2 v_i' \log(N\tilde{s})}}{2v_i'},$$

which implies that one root is positive and the other is negative. For the Desai-Fisher approximation (equation (16) in the main text) to hold, we must have $\tilde{s}/\tilde{U}_{bi} \gg 1$, i.e., $z_i$ must be positive. Therefore, we set

$$\tilde{U}_{bi} = \tilde{s} \exp\left[\frac{\tilde{s}^2 - \sqrt{\tilde{s}^4 + 8\tilde{s}^2 v_i' \log(N\tilde{s})}}{2v_i'}\right]. \tag{S7}$$

Now we turn to the choice of the function $\xi$. One simple class of functions that satisfy the constraints (S3), (S4) are the weighted averages

$$\tilde{s} = \xi(s_1, U_{b1}, s_2, U_{b2}, N) = \sum_{i=1}^{2} w_i\, s_i, \tag{S8}$$

with $w_1 \equiv w(s_1, U_{b1}, s_2, U_{b2}, N)$ and $w_2 \equiv w(s_2, U_{b2}, s_1, U_{b1}, N)$ and $w_1 + w_2 = 1$. In principle, the weight function $w$ can be any function of the population genetic parameters $s_1, U_{b1}, s_2, U_{b2}, N$ as long as $w_1 + w_2 = 1$. We tested three choices of these weights, (i) $w_i = 0.5$; (ii) $w_i = U_{bi}/U_b$; (iii) $w_i = s_i/(s_1 + s_2)$. To evaluate their accuracy, we plotted the rate of adaptation estimated empirically from our simulations against the theoretical values calculated using expression (S2) with $\tilde{s}$ given by equation (S8) and $\tilde{U}_{bi}$ given by expression (S7). We found that all weight choices show similarly good accuracy (Figure S2) and we chose the weights $w_i = s_i/(s_1 + s_2)$, such that

$$\tilde{s} = \sum_{i=1}^{2} \frac{s_i^2}{s_1 + s_2}. \tag{S9}$$

Taking these results together, we approximate the rate of improvement of trait $i$ in a two-trait organism as

$$r_i = \frac{\delta}{s_i} \times \begin{cases} 0 & \text{if } \frac{v_i'}{v_{\bar{i}}'} < 1/D, \\ \frac{\tilde{U}_{bi}}{\tilde{U}_b} f_{\mathrm{DF}}(\tilde{s}, \tilde{U}_b, N) & \text{if } 1/D \le \frac{v_i'}{v_{\bar{i}}'} \le D, \\ f_{\mathrm{DF}}(s_i, U_{bi}, N) & \text{if } \frac{v_i'}{v_{\bar{i}}'} > D, \end{cases} \tag{S10}$$

where $\bar{i}$ denotes the trait other than $i$ and $D$ denotes the ratio threshold. For all our subsequent calculations, we set the ratio threshold $D = 100$, but the specific choice of $D$ is inconsequential as long as $D \gg 1$ (Figure S3). To obtain evolutionary trajectories in the trait space, we substitute expression (S10) into equation (7) in the main text and numerically solve it using MATLAB function `ode45`.

### 3 Sampling genotypes with specific trait values in the pleiotropic GPFMs

In this section, we provide a justification for the procedure that we use to draw initial genotypes with a given trait value for our simulations on pleiotropic GPFMs (Section 2.4.4 in the main text). The challenge is that some trait values are represented by a very small fraction of genotypes, and obtaining such genotypes in uniform random samples is very difficult. This is particularly true for the pleiotropic GPFM with a low-entropy optimum. In principle, given the additive nature of our traits, it is possible to arrive at genotypes with the desired trait values by a biased random walk starting at the optimum. However, genotypes sampled in this way could be highly biased (e.g., they may contain unusually small number of sites with allele 1). A more principled way is to use the maximum entropy distribution under constraints (Bialek, 2012).

For any parameter $\boldsymbol{\lambda} = (\lambda_1, \lambda_2)^T$, we define the "tilted" distribution over the genotype space as

$$P(g; \boldsymbol{\lambda}) = \frac{\exp \langle \boldsymbol{\lambda}, \boldsymbol{y}(g) \rangle}{Z(\boldsymbol{\lambda})},$$

where for simplicity we defined the scaled trait value as

$$\boldsymbol{y}(g) = -\frac{\boldsymbol{x}(g)}{\delta} = \sum_{\ell=1}^{2L} g_\ell \boldsymbol{e}_\ell,$$
$$\boldsymbol{e}_\ell = -\boldsymbol{u}_\ell = -(\cos\theta_\ell, \sin\theta_\ell)$$

and $Z(\boldsymbol{\lambda})$ is the normalization constant. Since the probability $P(g; \boldsymbol{\lambda})$ is the same for all genotypes with the same traits $\boldsymbol{y}$, it is a maximum-entropy distribution for any given trait value. Importantly, $Z(\boldsymbol{\lambda})$ factorizes as

$$Z(\boldsymbol{\lambda}) = \prod_{\ell=1}^{2L} \left[1 + \exp\langle \boldsymbol{\lambda}, \boldsymbol{e}_\ell \rangle\right],$$

which implies that $P(g; \boldsymbol{\lambda})$ also factorizes as

$$P(g; \boldsymbol{\lambda}) = \prod_{\ell=1}^{2L} \frac{\exp\left[g_\ell \langle \boldsymbol{\lambda}, \boldsymbol{e}_\ell \rangle\right]}{1 + \exp\langle \boldsymbol{\lambda}, \boldsymbol{e}_\ell \rangle}.$$

In other words, under this tilted distribution, each site is in state 1 with probability

$$p_\ell(\boldsymbol{\lambda}) = \frac{\exp\langle \boldsymbol{\lambda}, \boldsymbol{e}_\ell \rangle}{1 + \exp\langle \boldsymbol{\lambda}, \boldsymbol{e}_\ell \rangle} \tag{S11}$$

and in state zero with probability $1 - p_\ell(\boldsymbol{\lambda})$. Re-writing equation (S11) in terms of the original site vectors $\boldsymbol{u}_\ell$, we have

$$p_\ell(\boldsymbol{\lambda}) = \frac{\exp\left[-\langle \boldsymbol{\lambda}, \boldsymbol{u}_\ell \rangle\right]}{1 + \exp\left[-\langle \boldsymbol{\lambda}, \boldsymbol{u}_\ell \rangle\right]} = \frac{1}{1 + \exp\langle \boldsymbol{\lambda}, \boldsymbol{u}_\ell \rangle}.$$

All of the above is true for any $\boldsymbol{\lambda}$. Finally, we set $\boldsymbol{\lambda}$ so that the expected trait value under this tilted distribution equals the desired trait value $\boldsymbol{x}_0$,

$$\mathbb{E}(\boldsymbol{x}; \boldsymbol{\lambda}) = \delta \sum_{\ell=1}^{2L} p_\ell(\boldsymbol{\lambda})\, \boldsymbol{u}_\ell = \boldsymbol{x}_0.$$

### 4 GPFMs with $M$ modules

In the main text, we considered GPFMs with just two functional modules (traits). While these models are tractable, their simplicity limits our ability to compare them with data. In biological organisms, such as *E. coli* evolving in the LTEE, the number of functional modules contributing to fitness (the so-called "phenotypic complexity") may be greater than two, although several studies suggest that for microbes this number may still be quite small, perhaps even below 10 (Kinsler et al., 2020; Iwasawa et al., 2022; Donyavi et al., 2025; Ghosh et al., 2026). Here, we simulate adaptation on the same four GPFMs as in the main text but with $M = 10$ functional modules/traits and ask whether the key features of the dynamics identified in the two-module models persist in this higher-dimensional trait space.

#### 4.1 Genotype-phenotype maps with $M$ modules

We use the same phenotype-fitness map defined by equation (1) in the main text, but now with $M = 10$. We consider the genome that contains $\Lambda$ sites (loci), partitioned into $M$ disjoint sets ("chromosomes") $\mathcal{C}_i$, with chromosome $\mathcal{C}_i$ containing $L_i$ loci, with $i = 1, \ldots, M$ and $\sum_i L_i = \Lambda$. All four genotype-phenotype maps can be written as

$$\boldsymbol{x}(g) = -\delta \sum_{\ell=1}^{\Lambda} g_\ell \, \boldsymbol{u}_\ell, \tag{S12}$$

where $\boldsymbol{u}_\ell$ is a unit vector associated with locus $\ell$; note that, unlike in equation (2) in the main text, allele 1 at locus $\ell$ moves the phenotype in the direction $-\boldsymbol{u}_\ell$. The differences between GPFMs arise from how $\boldsymbol{u}_\ell$ are determined, as described next.

**Pleiotropic GPFM with a high-entropy optimum.** Here, $\boldsymbol{u}_\ell$ are drawn randomly and independently from the unit sphere $S^{M-1}$ in an $M$-dimensional space. To obtain each $\boldsymbol{u}_\ell$, we draw a multivariate Gaussian random vector with a zero mean and a unit covariance matrix and normalize it to have unit length.

**Pleiotropic GPFM with a low-entropy optimum.** Here, $\boldsymbol{u}_\ell$ are drawn in the same way but restricted to the positive orthant. Specifically, we draw a Gaussian random vector as above, replace each of its components by its absolute value, and normalize it to unit length. Thus, $\boldsymbol{u}_\ell$ is uniformly distributed over the part of the unit sphere $S^{M-1}$ where all components are non-negative, and, according to equation (S12), allele 1 at any locus lowers all $M$ traits. For $M = 2$, this recovers exactly the pleiotropic GPFM with a low-entropy optimum analyzed in the main text.

**Modular GPFM with a high-entropy optimum.** Here, $\boldsymbol{u}_\ell = \pm \boldsymbol{e}_i$ for all $\ell \in \mathcal{C}_i$, where $\boldsymbol{e}_i$ is the unit vector along the $i$th axis. Exactly half of the loci on each chromosome have $\boldsymbol{u}_\ell = -\boldsymbol{e}_i$, so that allele 1 at these loci increases $x_i$, and the other half have $\boldsymbol{u}_\ell = +\boldsymbol{e}_i$. As in the two-module simulations, we simulate this GPFM by tracking only $x_i$.

**Modular GPFM with a low-entropy optimum.** Here, $\boldsymbol{u}_\ell = \boldsymbol{e}_i$ for all $\ell \in \mathcal{C}_i$ and $\boldsymbol{e}_i$ is a unit vector along the $i$th axis, which yields $x_i(g) = -\delta \sum_{\ell \in \mathcal{C}_i} g_\ell$, $i = 1, \ldots, M$. This GPFM is also simulated by tracking only $x_i$.

#### 4.2 Quantifying the distribution of adaptation across modules

With many modules, visualizing their performances and their rates of improvement becomes more challenging. We therefore developed two summary statistics to characterize these patterns. The first summary statistic—which we refer to as the "effective number of lagging modules"—captures the variation in how far from their optima different functional modules are. We define it as

$$n_{\text{lag}} = \left( \sum_{i=1}^{M} q_i^2 \right)^{-1}, \tag{S13}$$

where

$$q_i = \frac{|x_i|}{\sum_j |x_j|}$$

is the relative distance of trait $i$ to its optimum compared to other traits. If one module is much farther from its optimum than the others, then $n_{\text{lag}} \approx 1$. In contrast, if all modules are equally close to their optima, corresponding to the diagonal of the trait space, then $n_{\text{lag}} = M$. For $M = 2$, values of $n_{\text{lag}}$ near the maximum therefore correspond to a module performance ratio $R$ near 1 when both traits are negative, with $n_{\text{lag}} = \frac{(1+R)^2}{1+R^2}$.

The second summary statistic describes the variability in the rates of improvement across functional modules. For a window of $\Delta t$ generations, we calculate the improvement of each module $i$ as

$$I_i = \frac{\max\{x_i(t + \Delta t) - x_i(t), 0\}}{\delta}.$$

To make the analysis of our simulations analogous to that of the LTEE data in the main text, we use windows of $\Delta t = 10^4/6 \approx 1666.67$ generations, i.e., one-sixth of the simulated time course, the same fraction as the $10^4$-generation windows over the $6\times10^4$ generations of the LTEE data. Successive windows are shifted by $\Delta t/4 \approx 416.67$ generations, beginning at generation 1. At each window boundary we select the last recorded generation at or before that boundary, so that the intervals used in the calculation span 1666 or 1667 generations. Negative changes are set to zero separately in each replicate before pooling. Dividing each $I_i$ by the duration of the same window gives the same relative improvements $p_i$ defined below, because this factor cancels in the ratio. We calculate the corresponding relative improvement as

$$p_i = \frac{I_i}{\sum_j I_j}.$$

We define the effective number of modules targeted by selection as

$$n_{\text{sel}} = \left( \sum_{i=1}^{M} p_i^2 \right)^{-1}. \tag{S14}$$

If a single module improves and all others are stalled, then $n_{\text{sel}} \approx 1$. In contrast, if all modules improve by the same amount, then $n_{\text{sel}} = M$. Thus, larger values of $n_{\text{sel}}$ correspond to more modules improving simultaneously. Note that the summary statistic $n_{\text{sel}}$ is analogous to the effective number of selected genes that we use in our analysis of the LTEE data (Section 2.5 in "Methods"), where Simpson's index is calculated from the distribution of observed mutations among genes.

To make the simulation and LTEE analyses comparable, we pool replicate simulations within each time window (windows overlap by three quarters of their width, as in the LTEE analysis), just as mutations are pooled across the six LTEE populations. Because $I_i$ is calculated from changes in population mean phenotypes and need not be an integer, the pooled values are rounded to the nearest integers, so that each window yields, for each module, an integer number of improvement steps of size $\delta$. We refer to these steps as observations. Simpson's index depends on how many observations it is computed from, and since the number of observations differs between windows, we subsample each window to $K = 104$ observations (the number we used in the LTEE analysis). Specifically, we draw 104 observations without replacement and calculate Simpson's index $D_{\mathrm{S}} = \sum_i p_i^2$ from them. We repeat this subsampling $2 \times 10^3$ times, average $D_{\mathrm{S}}$ across subsamples, and report $n_{\mathrm{sel}} = 1/\overline{D_{\mathrm{S}}}$. The shaded bands in Figure S6 span the reciprocals of the 97.5th and 2.5th percentiles of $D_{\mathrm{S}}$ across subsamples. Windows containing fewer than $K$ observations are omitted.

#### 4.3 Simulations

##### 4.3.1 Regime and parameters

We simulated evolution on all four GPFMs in the concurrent mutations regime with complete linkage, with $\delta = 0.1$, $N = 10^4$ and $\mu = 10^{-5}$ per locus per generation. Strengths of selection were equal across modules ($a_i = 2\sqrt{2}$ for all $i$). In both pleiotropic GPFMs and in the modular GPFM with a low-entropy optimum, the genome contained $\Lambda = 400$ loci ($L_i = 40$ per module in the modular GPFM). In the modular GPFM with a high-entropy optimum, trait values are typically confined to the interval $[-L_i\delta/2, L_i\delta/2]$ from its optimum. Since for $L_i = 40$, the extreme trait values are only about $\pm 2$, which is less than the largest initial distance (about 3.0), we increased $L_i$ to 80 loci per module ($\Lambda = 800$).

##### 4.3.2 Initialization

**Choice of target initial trait coordinates.** We initialize populations at three locations in the trait space with different initial values of $n_{\mathrm{lag}}$ (Section 4.2), denoted by rhombi, upward-pointing triangles and downward-pointing triangles in Figure S6. For these three initial conditions, we define the weights

$$\boldsymbol{\omega}_1 = (c_1, 1, \ldots, 1)\,,$$
$$\boldsymbol{\omega}_2 = (c_2, c_2, c_2, 1, \ldots, 1)\,,$$
$$\boldsymbol{\omega}_3 = (1, 1, \ldots, 1)\,,$$

with $c_1 = 1/0.15 \approx 6.67$ and $c_2 = 4$ and set the target initial trait values at

$$\boldsymbol{x}_j^{\mathrm{tar}} = -d_0 \frac{\boldsymbol{\omega}_j}{\|\boldsymbol{\omega}_j\|},$$

where $d_0$ is chosen so that $\boldsymbol{x}_j^{\mathrm{tar}}$ lies on the fitness contour $W_0 = 0.25$, or equivalently $F_0 = \log W_0 = -\log 4 \approx -1.39$. Thus, at the initial condition 1, one trait is initially 6.67 times farther from its optimum than the others; at the initial condition 2, three traits are initially 4 times farther from their optima than the others; and at initial condition 3, all modules are equally far from their optima. These correspond to three target values of $n_{\mathrm{lag}} = 4.6$, 6.6 and 10.0.

**Choice initial genotypes on pleiotropic GPFMs.** We construct a genotype whose phenotype lies close to $\boldsymbol{x}_j^{\mathrm{tar}}$ in two steps; the procedure is the same for both pleiotropic GPFMs, which differ only in the directions $\boldsymbol{u}_\ell$. First, starting from the all-zero genotype, we go through the loci in order and switch a locus to allele 1 whenever this reduces the distance to $\boldsymbol{x}_j^{\mathrm{tar}}$. The resulting genotype is typically close to the target, but it is atypical, because it is assembled from the loci that come first in the list. Second, to obtain a more typical genotype at the same location, we place the same number of alleles 1 at randomly chosen loci and repeatedly swap the alleles at a randomly chosen locus carrying allele 1 and a randomly chosen locus carrying allele 0, accepting a swap whenever it reduces the distance to $\boldsymbol{x}_j^{\mathrm{tar}}$, until this distance falls below $\delta/2 = 0.05$ or $2.5 \times 10^5$ swaps have been attempted. If the resulting genotype is farther from the target than the one obtained in the first step, we use the latter. This procedure does not sample uniformly among genotypes near the target, unlike the maximum-entropy procedure used in the two-module simulations.

On the pleiotropic GPFM with a high-entropy optimum, this procedure yields initial genotypes with $n_{\mathrm{lag}}(0) = 4.6$, 6.6 and 10.0, i.e., on target. On the pleiotropic GPFM with a low-entropy optimum, this procedure yields initial genotypes with $n_{\mathrm{lag}}(0) = 8.2$, 8.5 and 10.0, i.e., the two imbalanced targets cannot be reached. This is because, with only $\Lambda = 400$ loci, there are very few sites whose vectors are nearly co-linear to any of the orthogonal basis vectors. All replicate populations for a given initial condition start from the same monomorphic genotype.

**Choice of initial trait values on modular GPFMs.** As in the two-module simulations (Section 2.4.4 in the main text), the initial phenotype is obtained by rounding each coordinate of $\boldsymbol{x}_j^{\mathrm{tar}}$ to the nearest multiple of $\delta$. This rounding shifts $n_{\mathrm{lag}}(0)$ by about 0.4, to 5.0, 6.2 and 10.0 on both modular GPFMs. All replicate populations for a given initial condition start from the same monomorphic trait vector.

##### 4.3.3 Termination and visualization

Each simulation ran for $10^4$ generations, with 30 replicate populations per initial condition. For $n_{\mathrm{lag}}$, we truncate each replicate trajectory once its mean fitness reaches $\overline{W} \geq 0.99$, the criterion used to terminate the two-module simulations. We also evaluate $n_{\mathrm{lag}}$ only while the total distance from the optimum satisfies $\sum_i |x_i| \geq 10\delta = 1$; closer to the optimum, each $|x_i|$ is only a few multiples of $\delta$, and $n_{\rm lag}$ jumps between coarse values. At each time point, we average $n_{\rm lag}$ over the replicate populations that satisfy both conditions and, as in Figures 2–4 in the main text, display this average only while at least 80% of the replicates (24 of 30) contribute. The statistic $n_{\rm sel}$ is computed from all replicates over the entire run; windows with fewer than $K$ pooled observations are omitted (Section 4.2).

##### 4.4 Results

The ten-module simulations reproduce the difference between modular and pleiotropic GPFMs that we found with two modules (Figure S6). On the modular GPFM with a low-entropy optimum, adaptation initially proceeds through only some of the modules that are away from their optima. In the first window, the most concentrated initial condition has $n_{\rm sel} \approx 1.3$ although $n_{\rm lag}(0) = 5.0$, and the intermediate condition has $n_{\rm sel} \approx 3.2$ with $n_{\rm lag}(0) = 6.2$ (Figure S6F,H). Figure S7 shows this in a single population, where the most strongly lagging module improves while the other nine change little. On the modular GPFM with a high-entropy optimum, adaptation is also concentrated at first, but less so, with $n_{\rm sel} \approx 3.0$ and 4.7 in the first window for the same two initial conditions (Figure S6B,D). On both modular GPFMs, $n_{\rm sel}$ then rises to 9.3–9.7 and catches up with $n_{\rm lag}$, and populations that started from different initial conditions reach similar, high values of $n_{\rm lag}$. The concentration of adaptation in a few modules is thus transient, and adaptation spreads to the remaining modules as the population approaches module-selection balance.

On the pleiotropic GPFM with a high-entropy optimum, $n_{\rm sel}$ is close to $n_{\rm lag}$ from the first window on (Figure S6A,C). In the first window, $n_{\rm sel} \approx 4.8$ and $n_{\rm lag} \approx 4.2$ for the most concentrated initial condition, and $n_{\rm sel} \approx 6.3$ and $n_{\rm lag} \approx 6.4$ for the intermediate one. On the pleiotropic GPFM with a low-entropy optimum, all three initial conditions start close to balance because the imbalanced targets cannot be reached (Section 4.3), and $n_{\rm sel}$ ($\approx 7.7$ and 7.8 in the first window for the two imbalanced conditions) also stays close to $n_{\rm lag}$ (Figure S6E,G). On both pleiotropic GPFMs, $n_{\rm sel}$ stays within about one module of $n_{\rm lag}$ throughout, so there is no period in which only a few of the lagging modules improve. With ten modules, as with two, adaptation on the modular GPFMs is first concentrated in a few modules and then spreads to all of them, whereas on the pleiotropic GPFMs it is spread across modules from the start. This bi-phasic pattern on the modular GPFMs is what motivates our interpretation of the LTEE data.

### 5 Supplementary Figures

**Figure S1. Numerical validation of results by Gomez et al. (2020) (next page). A.** Empirically estimated rates of adaptation in isolation $v_i'$ (solid circles) and the respective theoretical values based on equation (16) in the main text (open circles). Each data point corresponds to a unique combination of parameters $s_1$, $U_{b1}$, $s_2$, $U_{b2}$. Population size is $N = 10^4$. Points are color-coded by the ratio $v_2'/v_1'$, which takes values $1$ (black), $\approx 4.6$ (blue), $\approx 20.9$ (dark green), $\approx 95.6$ (purple), and $\geq 437$ (orange). The dashed line is the diagonal. **B.** Empirically estimated rates of adaptation in a two-module organism $v_i$. Colors correspond to panel A. Vertical dashed line represents a numerical cutoff for traits that improve very slowly. See Section 2.1 for details. **C.** Two-module rate $v_i$ (y-axis) plotted against the corresponding rate in isolation $v_i'$ (x-axis) in the "stalling regime" where $v_2'/v_1' > 100$. Crosses denote $v_1$; filled circles denote $v_2$. Colors correspond to panel A. Stalled traits (panel B) are omitted. The dashed line represents $v_i = v_i'$. **D.** Same as C but in the "simultaneous improvement regime" where $v_2'/v_1' \leq 100$. **E–F.** Ranges of $(s_i, U_{bi})$ corresponding to the parameter grid in panels A–D. For each theoretical pair $(v_1', v_2')$, four values of $s_i/U_{bi} \in \{10, 22, 46, 100\}$ are used to back-calculate $(s_i, U_{bi})$ per module, yielding $4 \times 4 = 16$ unique parameter combinations per grid point. Colors correspond to panel A.

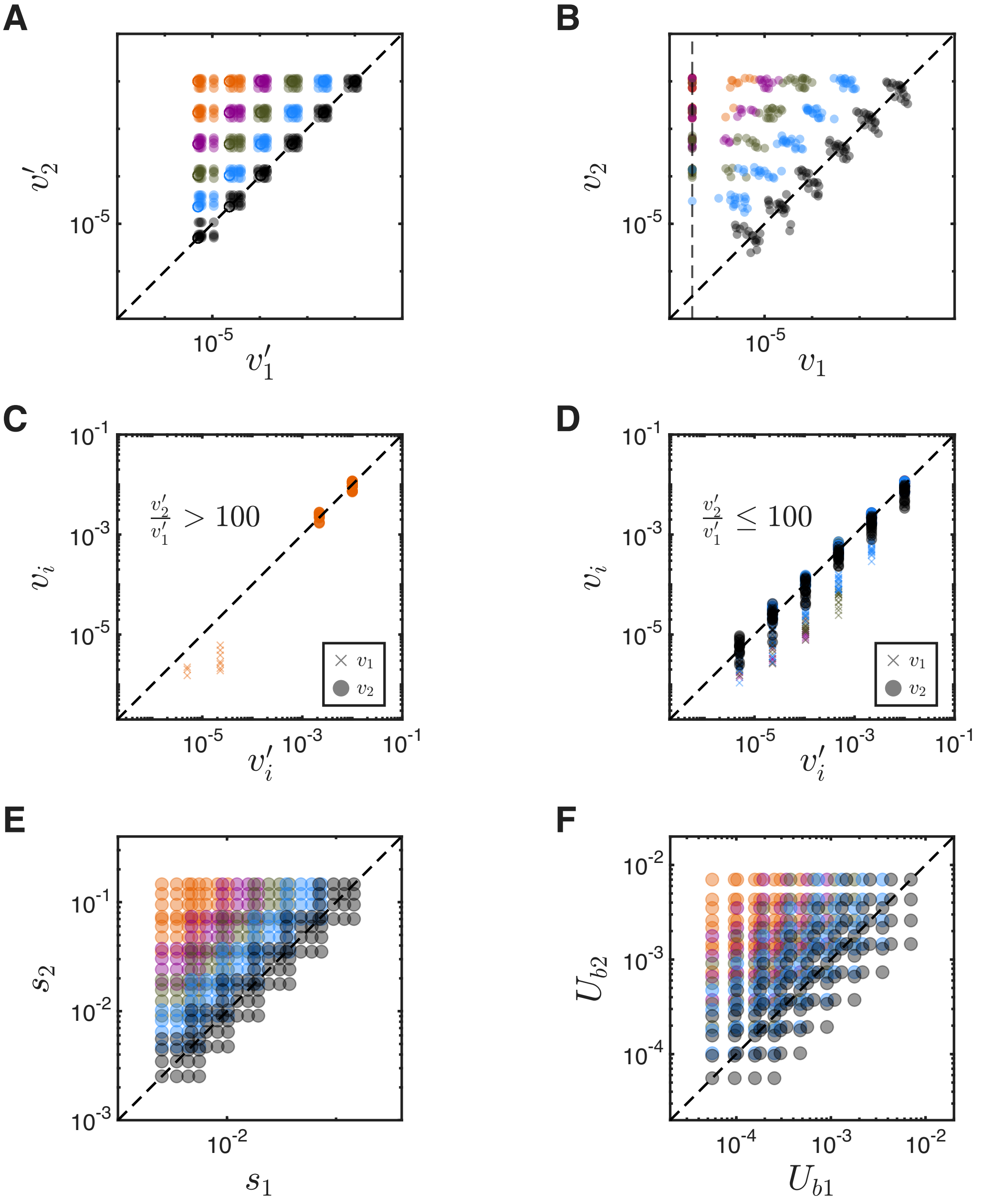
A
B
C
D
E
F
$v_2'$
$v_1'$
$v_2$
$v_1$
$v_i$
$v_i'$
$\frac{v_2'}{v_1'} > 100$
$\frac{v_2'}{v_1'} \leq 100$
$v_1$
$v_2$
$s_2$
$s_1$
$U_{b2}$
$U_{b1}$

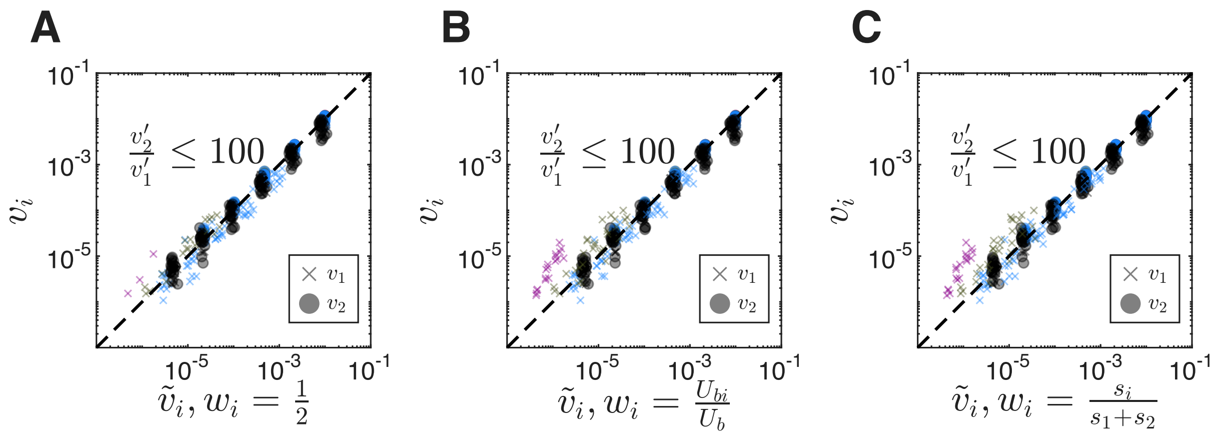

**Figure S2. Accuracy of the heuristic approximation for various choices of weights $w_i$.** We plot the empirically estimated rates of adaptation in a two-module organism $v_i$ against those predicted theoretically using equation (S2) in the simultaneous improvement regime ($v_2'/v_1' \leq 100$). Stalled traits (see Section 2.1) are omitted. We also omit parameter combinations for which the solution of equation (S6) has $\tilde{s}/\tilde{U}_{bi} < 10$, which places the effective two-trait organism outside the range of validity of the Desai–Fisher approximation. This is the case for 93, 54 and 45 of the 288 parameter combinations in panels A, B and C, respectively. Crosses denote $v_1$; filled circles denote $v_2$. Colors correspond to Figure S1A. Each panel corresponds to a different choice of weights $w_i$ for the calculation of the effective selection coefficient $\tilde{s}$ (equation (S8)), as indicated. The horizontal axis shows the $\tilde{v}_i$ predicted from equation (S2), evaluated with the weights specified in each panel, and the vertical-axis quantity $v_i$ is the empirically estimated rate. The dashed line represents $v_i = \tilde{v}_i$ in all panels.

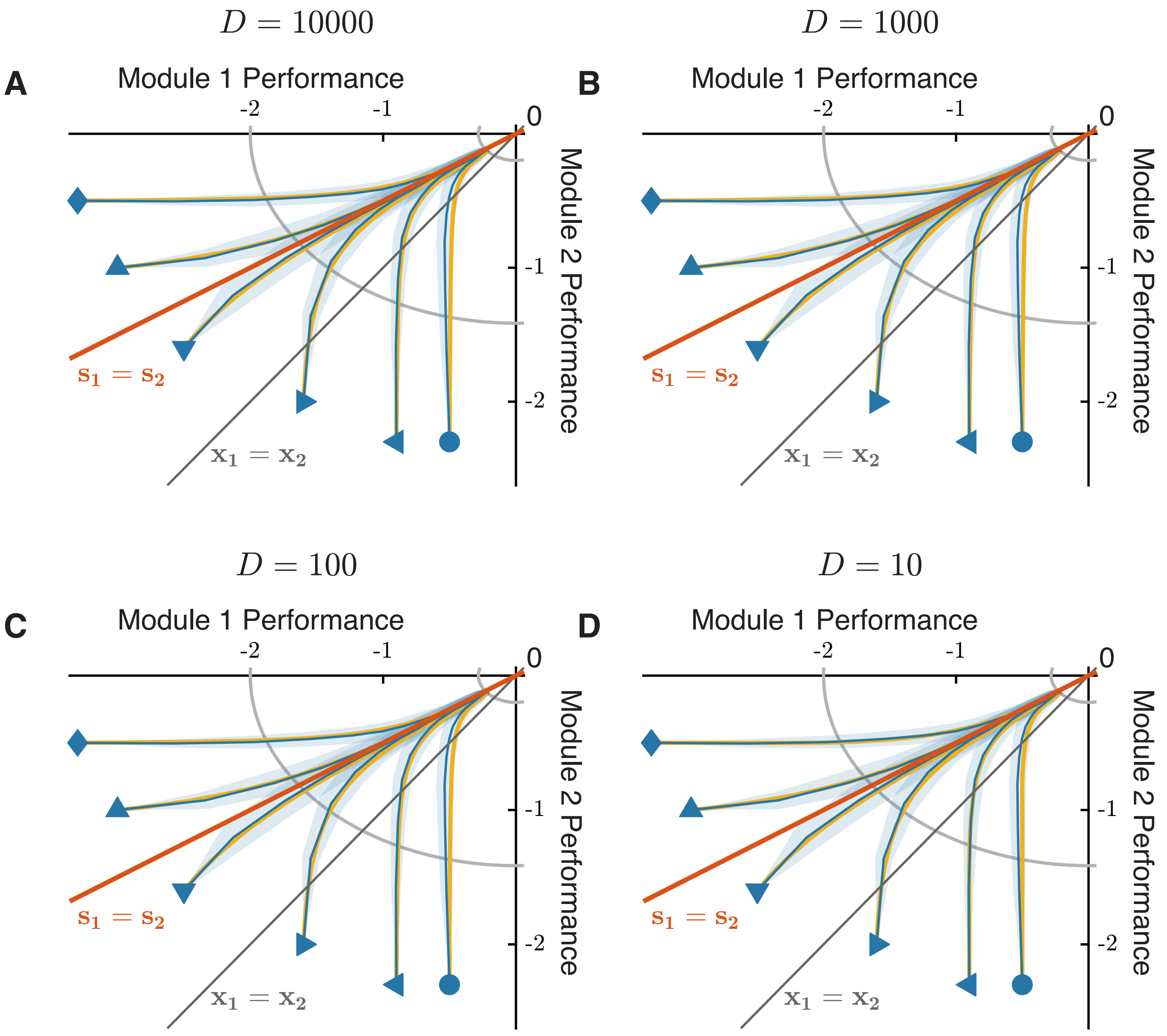

**Figure S3. Sensitivity analysis of the heuristic approximation with respect to the threshold parameter $D$.** Each panel shows the same simulated mean trajectories as Figure 3F (concurrent mutations, complete linkage; see main text for simulation details), alongside theoretical predictions (yellow) computed with a different value of $D$, as indicated.

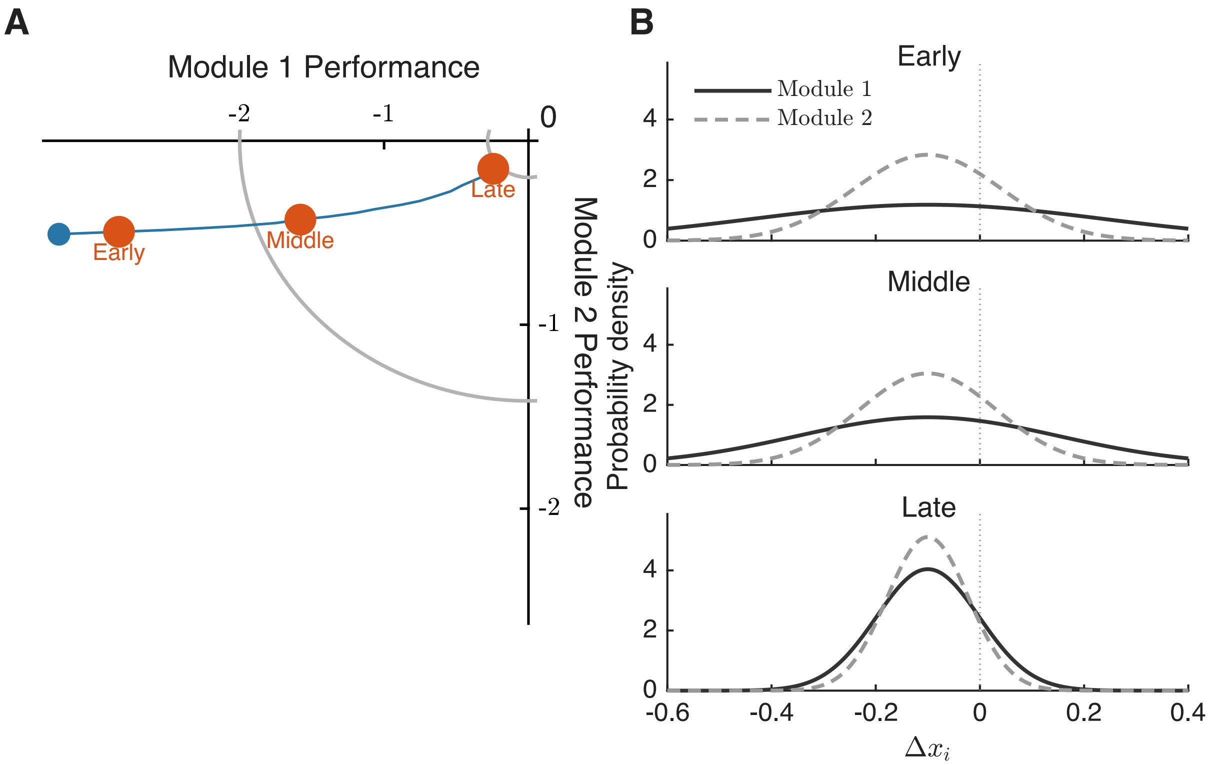

**Figure S4. Mutation-effect distributions in the nested Fisher's geometric model (nFGM) change along an adaptive path. A.** Mean evolutionary trajectory in the trait space for one initial condition of the nested FGM with $n_1 = n_2 = 10$ in the successive mutations regime (Figure 5 in the main text). Orange circles mark the points at 5%, 50% and 95% of the length of the trajectory, for which the distributions of mutational effects are shown in panel B. **B.** Distribution of the effects of mutations on trait 1 (solid) and trait 2 (dashed) at the three points marked in panel A.

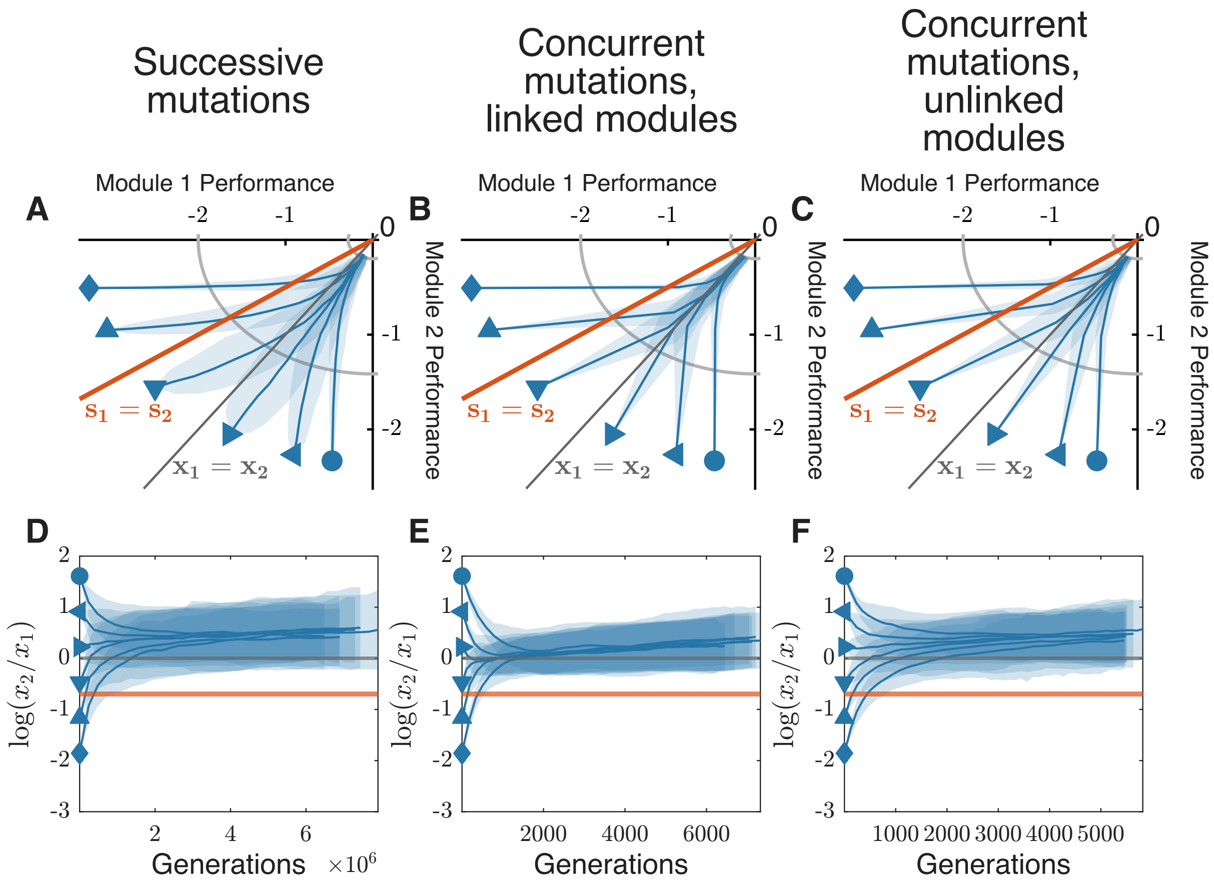

**Figure S5. Evolutionary dynamics in the nested FGM with different module dimensionalities.** Same as Figure 5D–I in the main text, but with $n_1 = 10$, $n_2 = 20$.

**Figure S6. Evolutionary dynamics on GPFMs with ten functional modules (next page).** All four GPFMs were simulated with $M = 10$ under concurrent mutations and complete linkage (Section 4.3). Left column, pleiotropic GPFMs; right column, modular GPFMs; top block, high-entropy optimum; bottom block, low-entropy optimum. Colors denote the three initial conditions, from the most concentrated (dark) to uniform (light), and markers indicate the first point of each curve. **A, B, E, F.** Effective number of lagging modules $n_{\text{lag}}$ (equation (S13)) over time, averaged over 30 replicates per initial condition. Shaded bands show $\pm 1$ standard deviation across replicates. Each replicate is truncated when its mean fitness reaches $\overline{W} \geq 0.99$, and the average is shown only while at least $80\%$ of replicates remain. **C, D, G, H.** Effective number of modules targeted by selection, $n_{\text{sel}}$ (equation (S14)), computed from improvements pooled across the 30 replicates in each time window. Lines show $n_{\text{sel}}$ calculated from $2 \times 10^3$ subsamples of $K = 104$ improvements each, drawn without replacement (Section 4.2), and shaded bands span the 2.5th to 97.5th percentiles across subsamples, as in the LTEE analysis (Figure 6). Each window is plotted at its midpoint, so the first point falls at generation $834$; windows with fewer than $K$ observations are not shown. Dashed and dotted horizontal lines mark $M = 10$ and $1$, respectively. On the pleiotropic GPFM with a low-entropy optimum (E, G), the two imbalanced initial conditions cannot be reached, so all three start close to balance (Section 4.3).

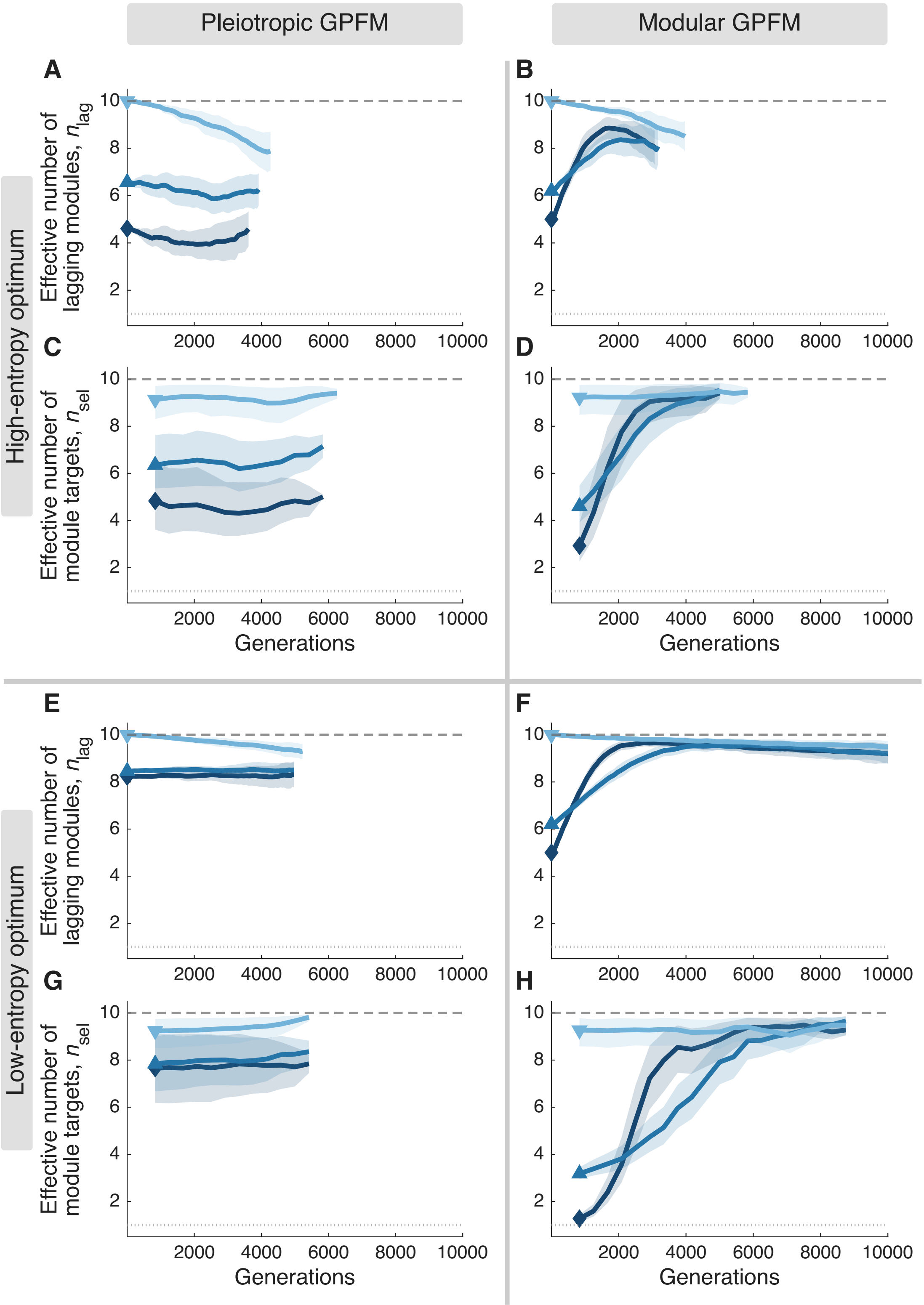

Pleiotropic GPFM
Modular GPFM
High-entropy optimum
Low-entropy optimum
A
B
C
D
E
F
G
H
Effective number of lagging modules, $n_{lag}$
Effective number of module targets, $n_{sel}$
Generations
2000 4000 6000 8000 10000
10 8 6 4 2

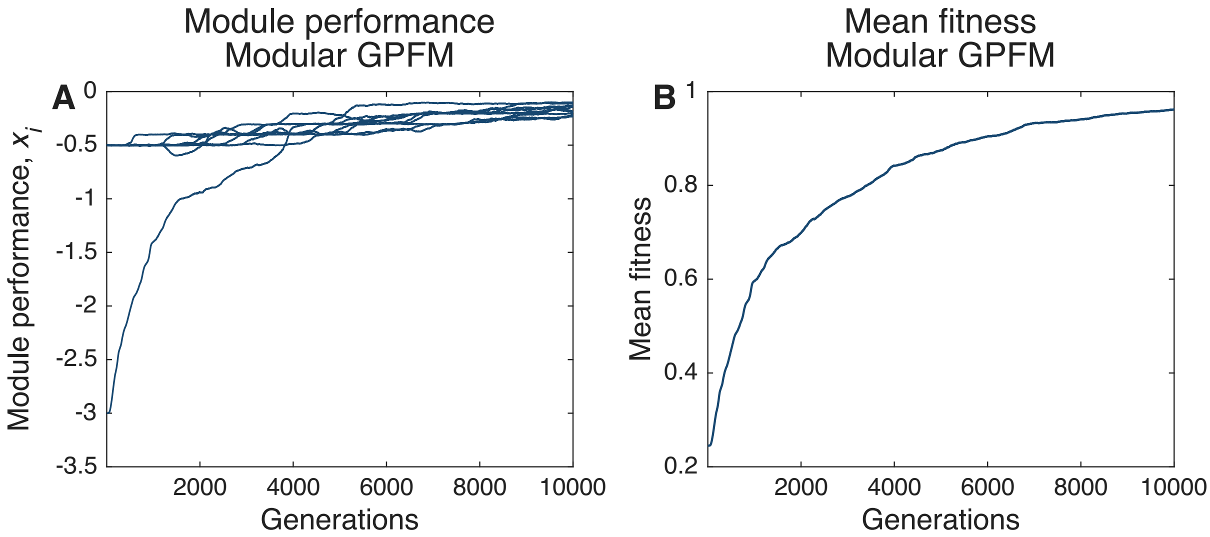

**Figure S7. Evolutionary stalling on the modular GPFM with a low-entropy optimum and ten functional modules. A.** Module performances $x_i$ over time for a single replicate initialized from the most concentrated initial condition (rhombus in Figure S6). Nine modules remain nearly stationary for the first $\sim 2000$ generations, while the most strongly lagging module improves from $-3.0$ to about $-1.0$. Thereafter, all ten modules improve together. **B.** Mean fitness for the same replicate. Parameters are given in Section 4.3.

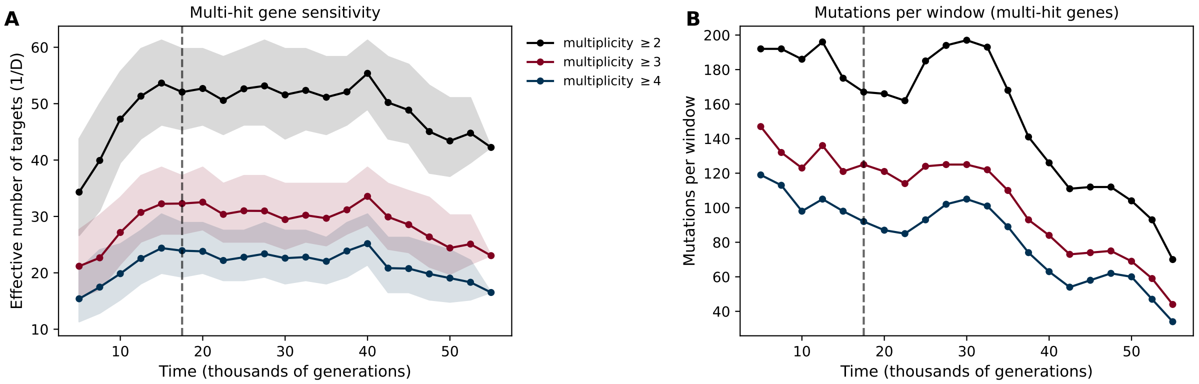

**Figure S8. Sensitivity of the sliding-window analysis to the multiplicity threshold. A.** Same as Figure 6 in the main text, but with mutations restricted to multi-hit genes with multiplicity thresholds greater than or equal to 2 (black; 262 genes, 902 mutations total, 70 subsampled per window), 3 (burgundy; 121 genes, 620 mutations total, 44 subsampled per window) or 4 (dark blue; 78 genes, 491 mutations total, 34 subsampled per window), as indicated in the legend. Slight declines at later time points are correlated with the overall decline in detections of mutations in multi-hit genes shown in panel B. **B.** Number of detected mutations per sliding window for each multiplicity threshold. Colors are the same as in panel A. Data from Good et al. (2017), six nonmutator LTEE populations.
